# *Toxoplasma* FER1 is a versatile and dynamic mediator of differential microneme trafficking and microneme exocytosis

**DOI:** 10.1101/2020.04.27.063628

**Authors:** Daniel N.A. Tagoe, Adeline Ribeiro E Silva, Allison A. Drozda, Isabelle Coppens, Bradley I. Coleman, Marc-Jan Gubbels

**Author notes:** Correspondence to Marc-Jan Gubbels. Daniel N.A. Tagoe and Adeline Ribeiro E Silva contributed equally to this work.

## Abstract

*Toxoplasma gondii* is a polarized cell concentrating several secretory organelles at the apical pole. The secretory micronemes come in two sub-populations differentiated by dependence on the Rab5A/C in their biogenesis. Calcium-dependent exocytosis of micronemes occurs at the very apical tip and is critical for parasite egress from its host cell, adhesion and invasion of the next cell. Ferlins represent a protein family with roles in exocytosis containing multiple Ca^2+^-sensing C2 domains. We determined that *T. gondii*’s ferlin 1 (FER1) localized dynamically to the parasite’s secretory pathway. FER1 function was dissected by dominant negative overexpression strategies. We demonstrated that FER1 traffics microneme organelles along the following trajectories:1. From the *trans*-Golgi-endosomes network to the subpellicular cortex; 2. Along the cortex to the apical end; 3. To the apical tip for fusion with the plasma membrane; 4. Retrograde transport allowing microneme recycling from mother to daughter; 5. Differential microneme sub-population traffic. Finally, FER1 overexpression triggers a microneme exocytosis burst, supporting the notion that the radially organized micronemes at the apical tip comprise a readily-releasable microneme pool. In summary, FER1 is pivotal for dynamic microneme trafficking, acts differently on the two microneme subpopulations, and acts on the plasma membrane fusion step during microneme exocytosis.

## Introduction

In humans, the apicomplexan parasite *Toxoplasma gondii* causes birth defects, vision loss, myocarditis and encephalitis. Lytic replication cycles unfolding through repetitive rounds of host cell invasion, intracellular replication and host cell egress are central to the pathogenesis of toxoplasmosis^1^. The micronemes are pivotal for successful host cell invasion as they contain adhesion molecules facilitating gliding motility and host cell attachment^2,3^. In addition, the *Toxoplasma gondii* micronemes encode a pore-forming protein, PLP1, that permeabilizes both the parasitophorous vacuole and host plasma membrane, which is required for host cell egress^4,5^. The parasite’s signal transduction pathways controlling the correct timing of micronemes secretion comprises cGMP, Ca^2+^ and phosphatidic acid (PA)^6^, accompanied by a crucial cAMP-mediated switch between the intracellular and extracellular states^7^.

The micronemes are localized along the apical cortex in association with the subpellicular microtubules emanating from the apical end^8,9^. Upon activation of exocytosis, micronemes move into the conoid, a tubulin basket at the apical tip of the parasites, to fuse with the plasma membrane. Following secretion, microneme protein complexes are embedded in the plasma membrane with the extracellular domains serving as adhesion domains and the cytoplasmic tail engaging with actin filaments that are transported in an apical to basal direction by a myosin motor^10^. A set of radial micronemes organized and anchored just below the conoid is believed to be a readily-releasable pool of micronemes^11–13^. Furthermore, microneme exocytosis is dosed to support prolonged periods of gliding motility enabling migration cross the biological barriers in between host cells^14^ as only ∼20% of all microneme proteins can be triggered to secrete at any given time^15,16^. Dosing is most likely achieved by gradually trafficking micronemes toward the conoid^15^. Finally, microneme exocytosis is balanced with active endocytosis^17^, whereas during cell division micronemes of the mother are re-directed into the forming daughters^18^.

Microneme biogenesis required protein trafficking through the secretory pathway comprising sequential passage through the endoplasmic reticulum (ER), Golgi apparatus, trans-Golgi network (TGN) and an endosome-like compartment (ELC)^19–21^. Microneme proteins undergo proteolytic processing to remove their pro-peptide inside the plant like vacuole (PLV or VAC), an acidic compartment^22–25^. Trafficking proteins from the ELC to micronemes and other destinations is facilitated by an unconventional endosomal system^20,21^. Protein sorting to both microneme and rhoptry secretory organelles requires sortilin (SORTLR) in the *trans*-Golgi^26^, trafficking to the rhoptries and dense granules REMIND^27^, whereas the HOPS/CORVET complex and Rab7 are involved in PLV/VAC routing^28^. Moreover, adaptor complex AP1 is involved in microneme and rhoptry protein trafficking but has a more general function across other vesicular trafficking events^29^. Although some rhoptry specific targeting signals have been identified^30,31^, specific sorting signals for microneme proteins are still elusive. Specific Rab GTPases have been associated with some aspects of microneme protein trafficking, and differentiate two sub-populations of micronemes with a different protein content: one of which is Rab5A/C-dependent, and one that is Rab5A/C-independent^9^. Consequently, specific microneme proteins end up in different, non-overlapping microneme sub-populations. To date, no differential function or behavior for these distinct microneme populations have been described, leaving the rationale for this distinction unresolved.

In all well-studied Ca^2+^ triggered exocytosis systems, proteins containing double C2 (DOC2) domains execute the Ca^2+^-mediated vesicle fusion^32^. The Apicomplexa encode an unconventional, long DOC2 domain protein essential for microneme exocytosis^33^. In addition, two ferlin proteins, FER1 and FER2, are conserved across the Apicomplexa^34,35^. FER1 in *Plasmodium berghei,* named ferlin-like protein (PbFLP), is reported to be essential for male gametocyte egress^36^, whereas *T. gondii* FER2 is required for rhoptry secretion^34^. The ferlins make up a unique branch of the DOC2 domain protein family because of their size (200-240 kDa) and contain five to seven C2 domains, organized in C2 pairs to form 2-3 DOC2 domains. The extended C2 repertoire in ferlins has broadened their functional spectrum beyond membrane fusion to vesicle trafficking and membrane repair^37^. Mammalian ferlins are differentiated by their sub-cellular localization at either the plasma membrane or on intracellular compartments, which relates to their function in late endosomal transit versus trans-Golgi recycling^38^. Mammalian otoferlin, essential for neurotransmitter release from the inner hair cells (IHC) in the auditory system, has been most widely studied. Otoferlin functions as both a scaffolding protein in the secretory pathway as well as in the actual membrane fusion during exocytosis^39–42^. Here we examined *T. gondii* FER1 and reveal roles in microneme protein trafficking, microneme organelle dynamics and exocytosis. Furthermore, FER1 acts differentially on the two different microneme populations.

## Results

### 1. *Toxoplasma* FER1 localizes to the secretory pathway

TgFER1 (TGGT1_309420) is a protein of 1425 amino acids harboring six C2 domains organized in three pairs and a tail-anchored transmembrane domain (**Fig. 1a**). A FerI domain found in most ferlins is present between the C2A and C2B domains, however no function has been associated with this domain^43^. In the genome wide fitness screen for the *Toxoplasma* lytic cycle, FER1 has a severe fitness score of -4.77, indicating that this protein is essential^44^. The HyperLOPIT sub-cellular localization atlas did not contain an assignment for TgFER1^45^. Hence, to localize FER1 we generated a polyclonal antiserum against the central C2DE domain, which is the most evolutionary diverse sequence of the protein compared to the other ferlins (**Fig. 1a**). By western blot, the α-FER1 serum reacts with the full-length FER1 protein with a predicted molecular weight of 159 kDa (**Fig. 1b**). Additional bands at approximately 120 and 30 kDa were detected, which most likely represent fragments of the full length FER1 protein, although cross reactivity with other proteins cannot be excluded. By IFA, since we anticipated a function in microneme exocytosis, we co-stained FER1 in both intracellular (**Fig. 1c**) and extracellular (**Fig. 1d**) parasites with MIC2 antiserum to highlight the micronemes. FER1 and MIC2 were physically distinct in intracellular parasites with FER1 localized apical of the nucleus. In extracellular parasites, FER1 localized throughout the cytoplasm, suggesting a re-localization in response to the environment. This coincides with the activation of microneme exocytosis and gliding motility^46^. The absence of FER1 co-localization with the micronemes suggests FER1 behaves like the mammalian ferlins in *trans*-Golgi recycling^38^.

**Figure 1.**
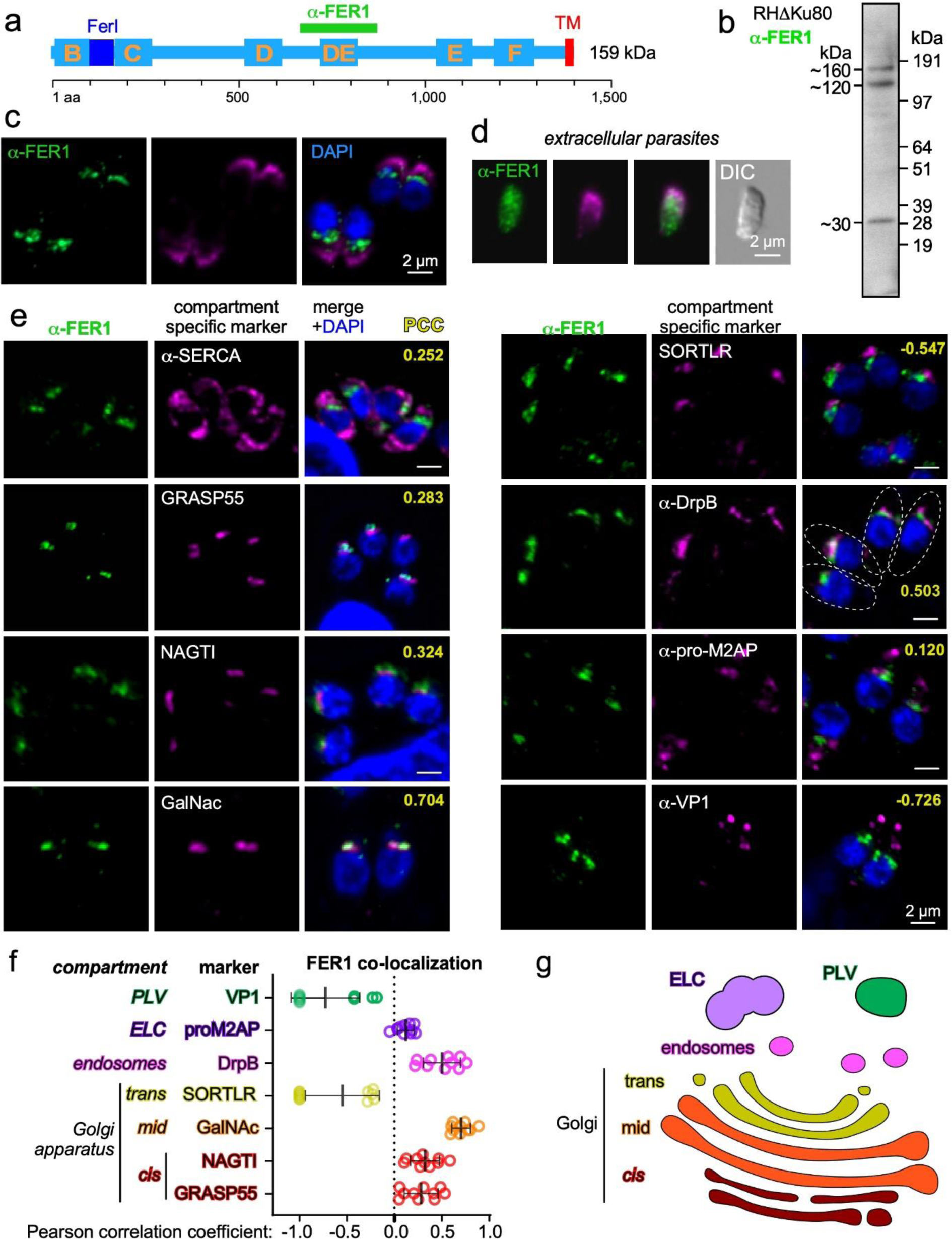
FER1 localizes to the secretory pathway. (**a**) Schematic representation of TgFER1. Yellow letters mark the C2 domains; the aa 669-877 region used to generate a specific antiserum is marked in green; FERI is a domain conserved in most ferlins with unknown function; TM is the transmembrane domain. (**b**) Cropped western blot analysis with the affinity purified guinea pig polyclonal antiserum generated against the FER1 region marked in panel A. Total lysate of wild type (RHΔKu80) parasites was loaded. (**c**) IFA of intracellular parasites and (**d**) extracellular parasites, co-stained for FER1 and MIC2 using respectively affinity purified FER1 antiserum and α-MIC2 serum. Samples were fixed with 4% PFA (**e**) Analysis of affinity purified FER1 antiserum localization in intracellularly replicating parasites by IFA co-stained with α-SERCA serum marking the endoplasmic reticulum (ER), GRASP55-RFP expressing parasites marking the *cis*-Golgi, NAGTI-YFP expressing parasites for the middle-Golgi, GalNac-YFP, and SORTLR-smOLLAS expressing parasites for the trans-Golgi, α-DrpB serum marking the *trans*-Golgi endosomes, α-Pro-M2AP serum marking the endosome like compartment (ELC) and, α-VP1 serum marking the plant like vacuole (PLV/VAC) and acidocalcisomes. PCC: Pearson correlation coefficients (yellow) for each image. (**f**) Pearson correlation coefficients reflect the level of FER1 colocalization with the compartment specific marker, n=10. (**g**) Schematic representation of FER1 localization through the secretory pathway. All IFA panels represent the wild type RHΔKu80 strain. See **Supplementary Figure S1** for marker cross-calibration.

To further define the sub-cellular compartment to which FER1 localizes in intracellular parasites we employed a series of *T. gondii* secretory pathway markers (**Fig. 1e-g**). We evaluated FER1 overlap with these markers using the Pearson correlation coefficient (PCC; ranging from 1 for perfect positive to -1 for perfect negative co-localization). Most of the FER1 signal overlaps with a marker for the *trans*-Golgi apparatus (GalNAc^29^) and the TGN-endosomal compartment (dynamin related protein B: DrpB^47)^. Surprisingly, no colocalization is observed with Sortilin, which is another marker for the *trans*-Golgi^26,47–51^. IFAs showing the relative distribution of the different markers in wild-type parasites are provided in **Supplementary Figure 1**. Although depletion of Sortilin does affect both microneme and rhoptry biogenesis^26^, it has a much stronger connection to sorting proteins to the rhoptries^19,29^. Thus, FER1 and sortilin could occupy different domains in the *trans*-Golgi, with the former catering to microneme and the latter to the rhoptry proteins. Taken together, in intracellular parasites FER1 localizes to the *trans*-Golgi-endosome network, whereas upon egress the signal disperses into the cytoplasm, supporting a role in the endosome-related trafficking pathway of microneme proteins.

### 2. Generation and validation of conditionally lethal FER1 mutant lines

To dissect the function of FER1 we attempted conditional depletion approaches by either placing the gene under a tetracycline regulatable promoter or fusing the auxin-inducible degron at either the N-or C-terminus. None of these efforts were successful, and neither were efforts to insert tags within the open reading frame, indicating that FER1 is extremely sensitive to perturbation. As an alternative, we resorted to a dominant negative (DN) approach. We generated an α-tubulin promoter driven N-terminal fusion of the destabilization domain (DD) linked to a Myc epitope that can be conditionally stabilized with Shield-1^52^. We established constructs without the 21 C-terminal amino acids encoding the TM domain (DD-Myc-FER1ΔTM) as well as a control construct encoding the full-length FER1 protein (DD-Myc-FER1^FL^) (**Fig. 2a**). We were able to generate stable parasite lines with both constructs in absence of Shield-1. Correct expression of the proteins was confirmed by western blot (**Fig. 2b**). We showed by plaque assay that overexpression of both the ΔTM and FL alleles caused parasite lethality (**Fig. 2c**). Collectively, these data align with a critical role of FER1 in the lytic cycle.

**Figure 2.**
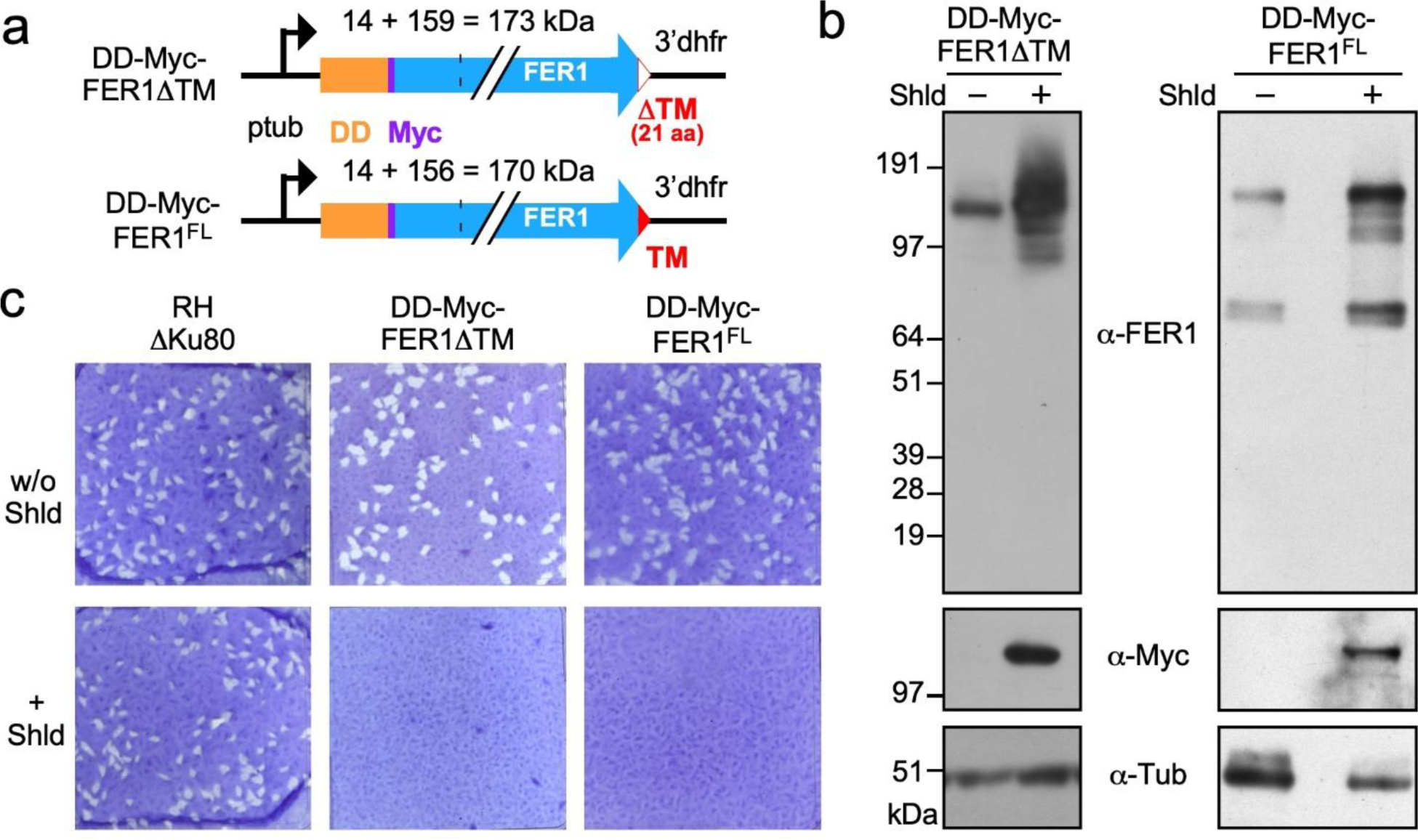
Generation and validation of conditional FER1 overexpression parasite lines. (**a**) Schematic representation of the overexpression constructs driven by the strong constitutive α-tubulin promoter (*ptub*). DD: destabilization domain; Myc: cMyc epitope tag; TM: transmembrane domain; FL: full length. (**b**) Western blot analysis of the overexpression parasite lines. Polyclonal guinea pig FER1 antiserum as in Fig 1. Monoclonal antibody12G10 recognizing α-tubulin was used as loading control. Parasites were induced with 1 µM Shield-1 for 24 hrs. (**c**) Plaque assays of infected HFF monolayers grown for 7 days ±1 µM Shield-1.

### 3. FER1ΔTM overexpression causes a microneme exocytosis defect

To determine the cause of the lethal defect upon FER1ΔTM overexpression, we evaluated key events in the lytic cycle. Host cell invasion was strongly reduced, consistent with an anticipated FER1 role in microneme exocytosis (**Fig. 3a**). We observed a mild reduction in the parasite multiplication rate, with a slight but significant (*p*=0.031) accumulation of 4-parasites/vacuole (**Fig. 3b**). However, no specific delay was observed in a particular cell cycle stage (**Fig. 3c**), suggesting that the overall rate through cell division is somewhat reduced. We triggered microneme-exocytosis mediated egress with Ca^2+^-ionophore A23187 and assessed the integrity of the parasitophorous vacuole membrane (PVM) using GRA3 as marker together with IMC3 marking the parasite’s cytoskeleton (**Fig. 3d**). We observed a 90% egress reduction in the overexpression condition, which was confirmed using pharmacological triggers in different points in the signaling pathway (**Suppl Fig. S2a,b**). Together, this convincingly supports a microneme defect (**Fig. 3e**). Trail assays were performed to further functionally assess microneme exocytosis and we determined that gliding motility is abrogated (**Fig. 3f**). Finally, we determined differential exocytosis of the Rab5A/C-dependent (MIC3, 5, 8) and -independent (MIC2, M2AP,etc.) microneme sub-populations^9^. The latter sub-population was tested by release of processed MIC2 in the medium under various triggers, whereas the former was visualized by MIC3, 5 and 8 protein exposure on the parasite’s surface. Overall, we see a sharp reduction in exocytosis to nearly undetectable levels of both microneme populations in the FER1ΔTM overexpressing mutant (**Fig. 3g,h**; **Suppl Fig. S2c**). However, we detected propranolol induced exocytosis at a 10-fold lower level than the non-induced control (**Fig. 3g**). Additionally, a small amount of MIC3 on the surface of induced parasites was observed, indicating a minimal exocytosis capacity (**Fig. 3h**). Since propranolol triggers the Ca^2+^-independent PA pathway, this section of the pathway appears to be still functional. Taken together, this suggests that FER1 acts in the Ca^2+^-dependent events in the microneme protein secretory pathway.

**Figure 3.**
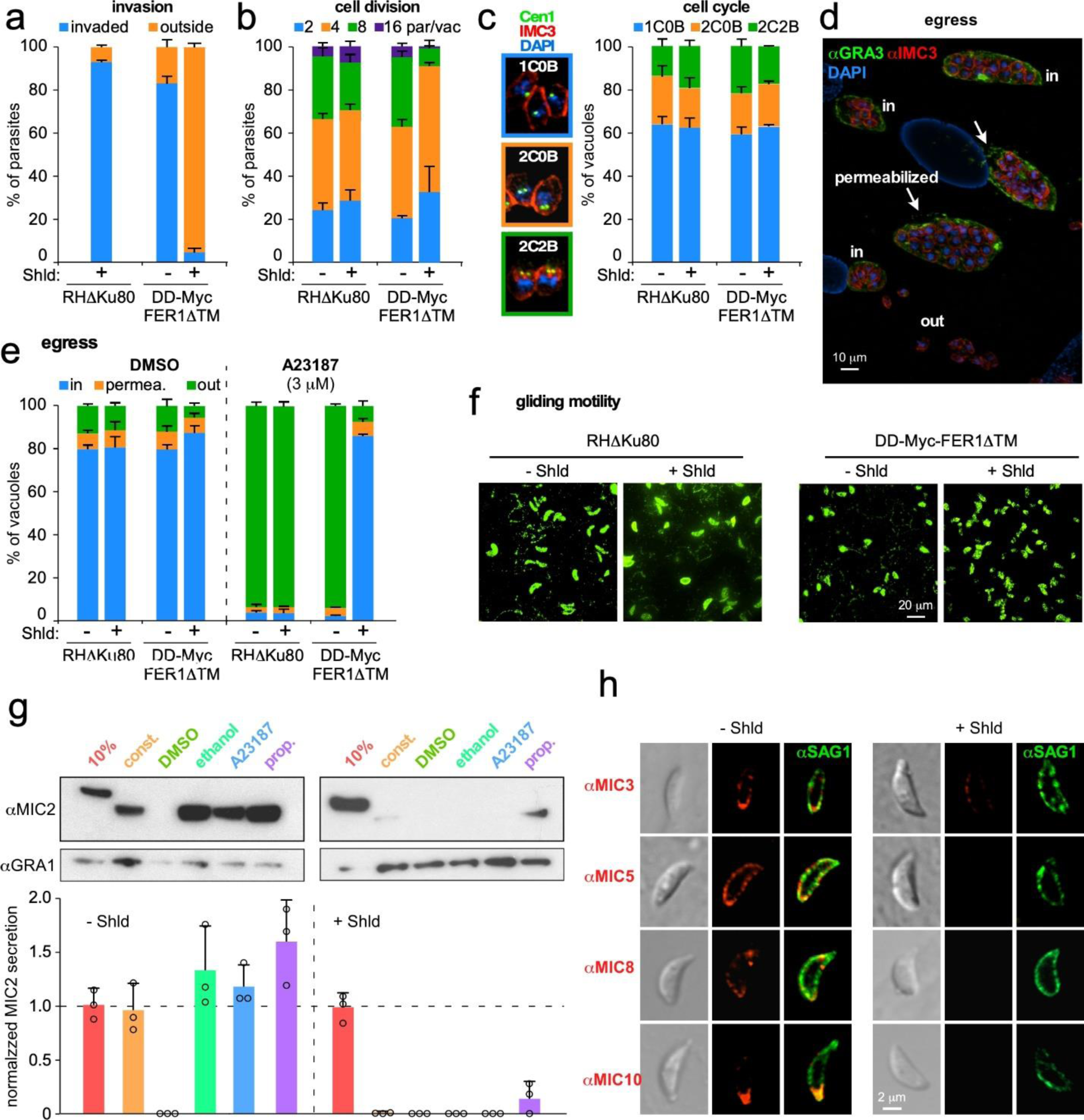
Phenotypic characterization of parasites overexpressing DD-Myc-FER1ΔTM. (**a**) Red-green invasion assay reveal an invasion defect. n=3+std. (**b, c**) Cell division and cell cycle progression analysis. Cell cycle stages quantified in the right of panel **c** are shown on the left. “C” and “B” represent the number of centrosomes and daughter buds, respectively. Parasites were allowed to invade for 2 hrs upon which 1 µM Shield-1 was added for 18 hrs. p=0.031 (*t*-test) accumulation of parasite in the 4-cells/vacuole stage. n=3+std. (**d,e**) Induced egress assays. Parasites grown for 30 hrs and induced for 18 hrs with 1 µM Shield-1 were triggered for egress with calcium ionophore A23187 (or vehicle control), fixed and stained with α-IMC3 (parasite cortex) and α-GRA3 (PVM) sera and scored for status of vacuole permeabilization and/or egress (**d**; representative image; arrows mark holes in the PVM of permeabilized vacuoles). n=3±std. Parasites were not under Shield-1 pressure during the 5 min pharmacological incubation. **(f)** Trail assay using α-SAG1 serum to assess gliding motility. Parasites were induced for 18 hrs ± 1 µM Shield-1, mechanically released from host cells and kept under 1 µM Shield-1 throughout the 30 min gliding experiment at 37°C. (**g**) Assessment of microneme secretion by western blot detection of MIC2 released in the supernatant under various triggers. Parasites were induced 18 hrs with 1 µM Shield-1 and harvested by physical release from the host cell. 10%: 10% of total lysate; const.: 1 hr constitutive secretion at 37°C (no secretagogue); 1% ethanol; 2 µM A23187; 500 µM propranolol. DMSO is the vehicle control for A23187. Induced secretion for 5 min at 37°C. Bottom of panel represents quantified secretion normalized to the GRA1 signal and to the 10% loading control for each condition. n=3+std. Parent line controls in **Supplementary Figure S2**. Parasites were not under Shield-1 pressure during secretion assay. (**h**) Secretion of the Rab5A/C-dependent microneme population was assessed by IFA on non-permeabilized parasites induced for 18 hrs ± 1 µM Shield-1, mechanically released, and exposed to fresh host cells for 5 min at 37°C. Parasites were not under Shield-1 pressure during assay.

### 4. FER1ΔTM overexpression causes a microneme trafficking defect

By design, we anticipated that FER1ΔTM would mislocalize. IFA in the DD-Myc-FER1ΔTM line revealed a reproducible signal accumulation in a defined mid-apical region upon Shield-1 induction (**Fig. 4a**). In parallel, we generated a YFP-tagged version of the FER1ΔTM allele to directly track accumulation. We observe no YFP signal without Shield-1 whereas under Shield-1 YFP accumulates in the mid-apical region as seen with the Myc construct, displaying a PCC of 0.968 co-localization with MIC2 (**Fig. 4b**; **Suppl Fig. S3a**). Co-staining with Rab5A/C-dependent (MIC5) class showed perfect overlap with the accumulated YFP signal indicating that all microneme proteins are misdirected (**Fig. 4c**; **Suppl Fig. S3b**). The α-FER1 signal also co-localizes with the YFP signal, suggesting that endogenous FER1 is also diverted from its *trans*-Golgi-endosome localization (**Fig. 4d**). Collectively, these data indicate that both the Rab5A/C-dependent and -independent microneme sub-populations are controlled by FER1.

**Figure 4.**
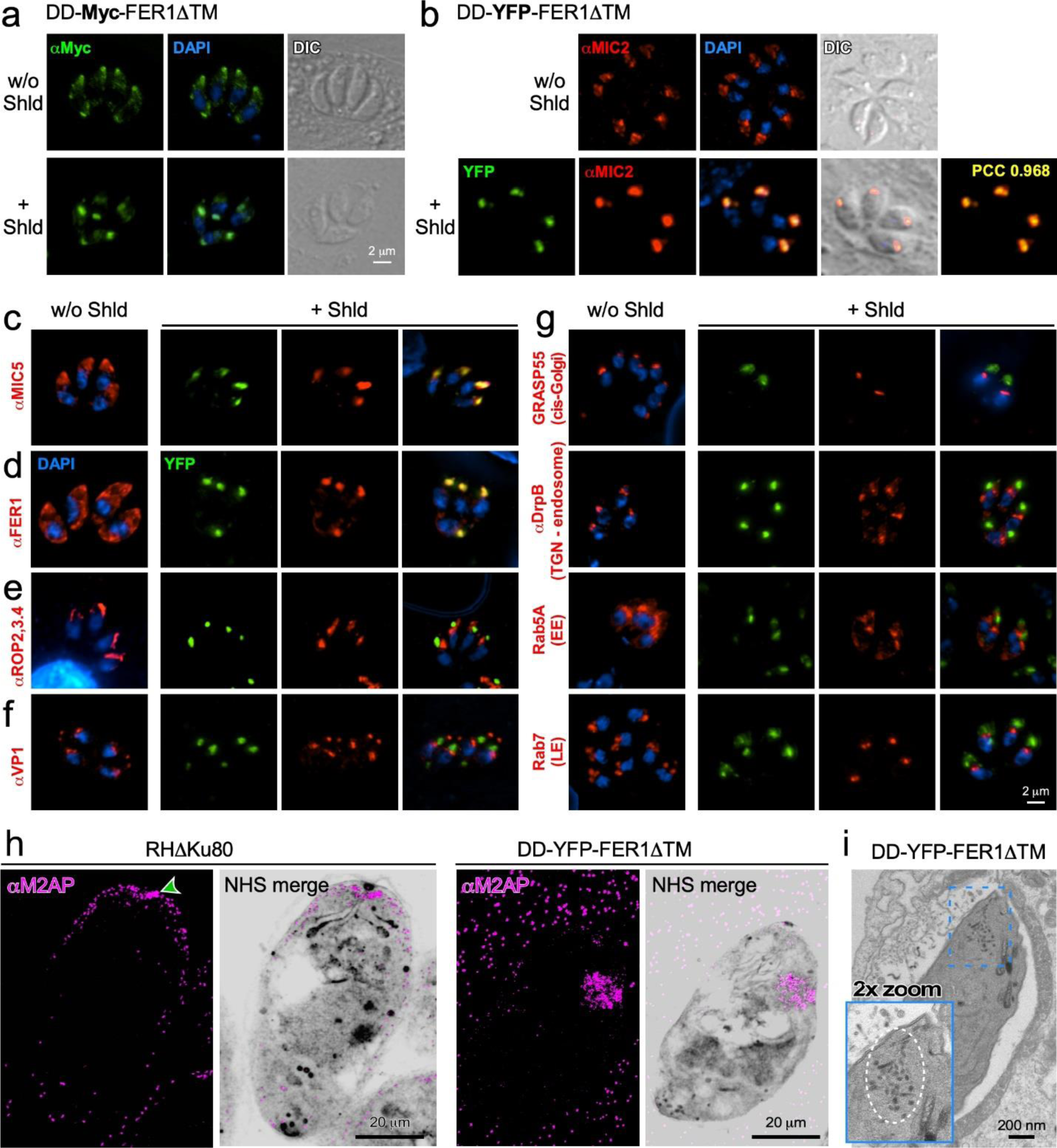
Overexpression of dominant negative FER1ΔTM constructs result in microneme mis-localization. **(a, b**) Overexpression of DD-Myc-FER1ΔTM (**a**) or DD-YFP-FER1ΔTM (**b**) leads to accumulation of microneme protein, visualized with α-MIC2, in a central, apical location. YFP and MIC2 co-localization was assesses by Pearson correlation coefficient. (**c**) Microneme proteins of the Rab5A/C-dependent trafficking pathway, visualized with α-MIC5, also accumulate in the FER1 compartment. (**d**) α-FER1 serum confirms exclusive accumulation in the microneme protein compartment. (**e**) ROP proteins do not accumulate and rhoptry morphology is normal. (**f**) Co-localization of VP1 and the YFP accumulation is not detected. (**g**) Markers for *cis-*Golgi, *trans*-Golgi network (TGN), as well as early (EE) and late (LE) endosome markers localize normally and do no co-localize with the YFP accumulation. In all IFA experiments parasites were treated with 1 µM Shield-1 for 18 hrs. (**h**) Pan-expansion microscopy of RHΔku80 parent line and DD-YFP-FER1ΔTM overexpressing parasites induced for 18 hrs with 1 µM Shield-1. Compiled stacks of 20 slices are shown. Green arrowhead marks the radial set of microneme proteins in the RHΔku80 parasites. (**i**) TEM of DD-Myc-FER1ΔTM overexpressing parasites induced for 16 hrs with 1 µM Shield-1. Dotted circle marks cytoplasmic microneme accumulation in the apical cytosol, which is enlarged in the inset.

Since all known micronemes trafficking mutants present defects in rhoptry protein trafficking^23^, we interrogated the rhoptries by IFA. ROP proteins remained localized to rhoptries, displaying their normal morphology and distribution (**Fig. 4e**). This distinction makes the DN-FER1ΔTM unique by disrupting microneme trafficking while leaving the rhoptries intact.

The mid-apical localization is reminiscent of the position of various compartments of the *Toxoplasma* secretory pathway. To differentiate whether the accumulation is due to a protein trafficking problem, or is of another nature, we again used a series of compartment-specific markers (**Fig. 4f, g**). No overlap was detected between the PLV/VAC (VP1) and FER1 (YFP) signals (**Fig. 4f**, **Suppl Fig. S3c**). Furthermore, no co-localization was observed with the Golgi or endosomal compartments. All markers (GRASP55, DrpB, Rab5A, Rab7) displayed their normal morphology (**Fig. 4g**). Instead, DN-FER1ΔTM accumulated in an uncharacterized compartment, beyond known trafficking steps toward the micronemes and, after splitting from rhoptry protein trafficking^23^.

To reveal the sites of FER1ΔTM and micronemes accumulation, we performed a two-step ultrastructure expansion microscopy (pan-ExM). We stained the micronemes with M2AP antiserum and as a guide co-stained general protein density with the NHS-ester. In parent parasites, micronemes are observed in both channels as protein-dense small, elongated foci aligned along the parasite’s cortex and a set radially arranged at the apical pole (**Fig. 4h**). In mutant parasites, both channels reveal mature micronemes mis-localized in the mid-apical region. Transmission electron microscopy (TEM: **Fig. 4i**) confirmed that micronemes with normal ultrastructure are absent from the cortex but aggregate in the parasite’s apical cytoplasm. Immune-electron microscopy (IEM) using MIC2 antiserum firmed up this assessment. Gold particles positioned MIC2 in the aggregated microneme structures (**Suppl Fig. S2d**). These could either be the result of retrograde trafficked mature micronemes released from the cortex, or, an incomplete organelle biogenesis and/or recycling of micronemes from mother to daughters during cell division.

In summary, overexpression of DN-FER1ΔTM leads to mis-localization of fully formed micronemes without disrupting the rhoptries, indicating that FER1 functions in the endosome-related trafficking and/or biogenesis of micronemes beyond the TGN^19^.

### 5. DD-YFP-FER1ΔTM overexpression retracts micronemes from the cortex

To differentiate between the retrograde trafficking from the biogenesis scenarios, we asked whether the aggregated microneme are proteolytically processed by removal of the pro-peptide in the ELC/PLV compartments^22,25^. For that, specific antisera against the pro-peptide of M2AP and the mature M2AP protein were used^53^. No difference in relative abundance of the pro-M2AP versus the total amount of M2AP protein was observed (**Fig. 5a**). Furthermore, by IFA we observed normal MIC5 pro-peptide containing proteins in the ELC compartment^53^, and no co-localization with the FER1 accumulation (**Fig. 5b**). These observations indicate that the mis-localized micronemes contain fully mature proteins.

**Figure 5.**
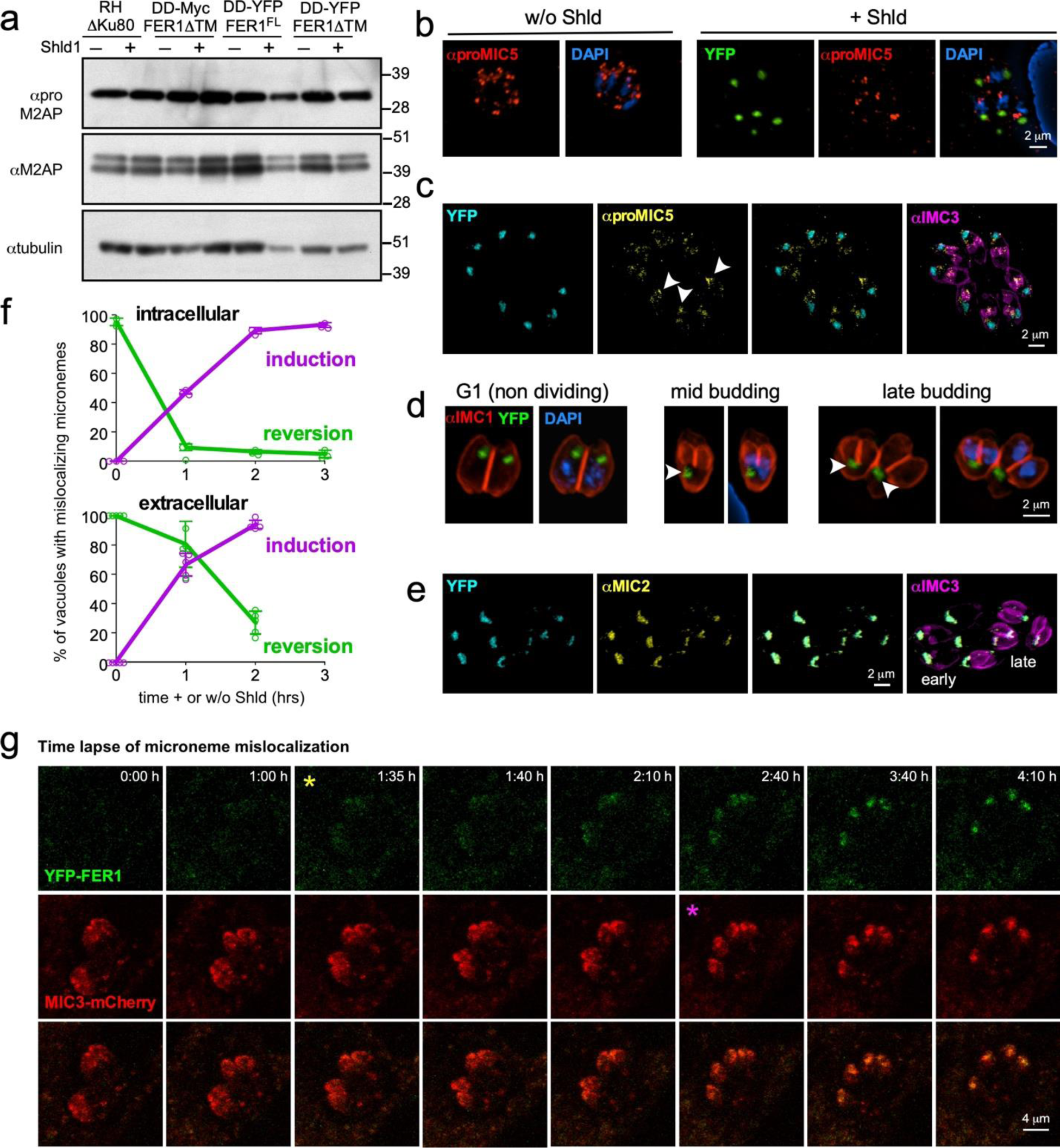
Microneme trafficking dynamics upon DD-YFP-FER1ΔTM overexpression. (**a**) Normal maturation by pro-peptide processing in the ELC/PLV compartments was demonstrated by western blot using serum against the M2AP pro-peptide (α-proM2AP) and the mature protein (α-M2AP). α-tubulin serves as loading control. (**b**) IFA showing the normal localization of proMIC5 in the ELC/PLV compartment in newly forming daughter buds using α-proMIC5. (**c**) Induced mutants co-stained with YFP, proMIC5, and IMC3 antiserum to track the timing and localization of proMIC5. Arrowheads mark the proMIC5 compartment within daughter buds. DD-YFP-FER1ΔTM is exclusively present in the mother parasites. (**d**) Induced mutants co-stained with IMC1 antiserum to track division stages as indicated. Arrowheads mark the mother parasite’s YFP accumulation migrating into a basal direction during progression of daughter budding. Note the absence of YFP in the daughter buds. (**e**) Induced mutants co-stained with YFP, MIC2, and IMC3 antisera to track the localization of mature MIC2 protein through cell division. DD-YFP-FER1ΔTM consistently colocalize with MIC2 throughout the division stages. (**f**) Time courses of the incidence of vacuoles or individual parasites with mis-localizing micronemes in intracellular and extracellular DD-YFP-FER1ΔTM parasites visualized through α-MIC2 IFA. At least 100 vacuoles (intracellular) or parasites (extracellular) were counted per time point in reversion experiments. n=3±std (intracellular) or n=5±std (extracellular). (**g**) Select panels from time lapse experiment with DD-YFP-FER1ΔTM parasites co-transfected with MIC3-mCherry to track microneme localization dynamics. At t=0, 2 μM Shield-1 was added. Yellow asterisk marks the first time point at which the piling up of YFP-FER1 could be convincingly observed; while purple asterisk marks the first time where MIC3-mCherry can be seen re-localizing from the apical cortex to the central apical localization co-localizing with YFP-FER1. Panels from supplementary movie S1. In all experiments parasites were treated with 1 µM Shield-1 for 18 hrs.

Since pro-MIC5 manifests predominantly in budding parasites, the co-staining with FER1 revealed another intriguing phenomenon: no DD-YFP-FER1ΔTM accumulates in newly forming parasites. We validated this by triple staining with YFP, α-pro-MIC5 and α-IMC3 (**Fig. 5c**). Moreover, representative images demonstrate that YFP migrates basally toward the residual body in parasites undergoing division (**Fig. 5d**). We never observed any new micronemes in the apical periphery of daughter buds either (**Fig. 5e**). Collectively, we observed that mature micronemes progress normally in forming daughters, yet new micronemes do not assemble in the daughters. This suggest that microneme proteins end up in the YFP-FER1ΔTM pile up formed in the mother and cannot be recycled to the daughter cells.

This raised the next intriguing question: is microneme aggregation independent on cell division? This model implies that micronemes can detach from the cortex and be directed toward the FER1ΔTM microneme aggregate. To this end we performed time courses to determine induction and reversion kinetics of the Shield-1 induced phenotype in both intracellular and extracellular parasites. Microneme mis-localization kinetics were over 95% complete within two hours (**Fig. 5f** top panel), which is much shorter than the 6.5 hr cell division cycle, indicating that the process is cell cycle independent^54^. The same kinetics were also observed in extracellular parasites which are arrested in a non-division G1a/0 state (**Fig. 5f** bottom panel)^55^. Finally, time lapse microscopy allowed us to visualize the cell division independent re-localization of the micronemes following Shield-1 induction. In DD-YFP-FER1ΔTM co-expressing a MIC3-mCherryRFP marker we observed that microneme retraction and accumulation with YFP-FER1ΔTM started after 1.5 hr of Shield-1 induction (**Supplementary Movie S1** and **Fig. 5g**). Moreover, we confirmed that microneme retraction was independent of daughter budding as tracked with a marker enriched in daughter cytoskeletons, IMC3-mCherryRFP (**Supplementary Movie S2** and **Suppl Fig. S3**).

Taken together, our results indicate that intact organelle microneme trafficking is cell cycle independent and bidirectional: either in apical direction to the site of exocytosis or, from the peripheral cortex to the site of aggregation that could be a relevant natural process when micronemes are recycled from mother to daughters^18^.

### 6. FER1^FL^ overexpression results in premature, untriggered microneme exocytosis

We dissected the phenotype associated to DD-Myc-FER1^FL^, by assessing the localization of FER1, which upon Shield-1 induction presents on membrane structures throughout the parasite. (**Fig. 6a**). We asked whether these parasites display changes in their microneme exocytosis ability using the same set of secretagogues applied previously (**Fig. 3g**). We observe strong exocytosis under all conditions (**Fig. 6b**). Although slight variations were observed in the sensitivity to different secretagogues (**Fig. 3g; Suppl Fig. S1c**), they were not significant.

**Figure 6.**
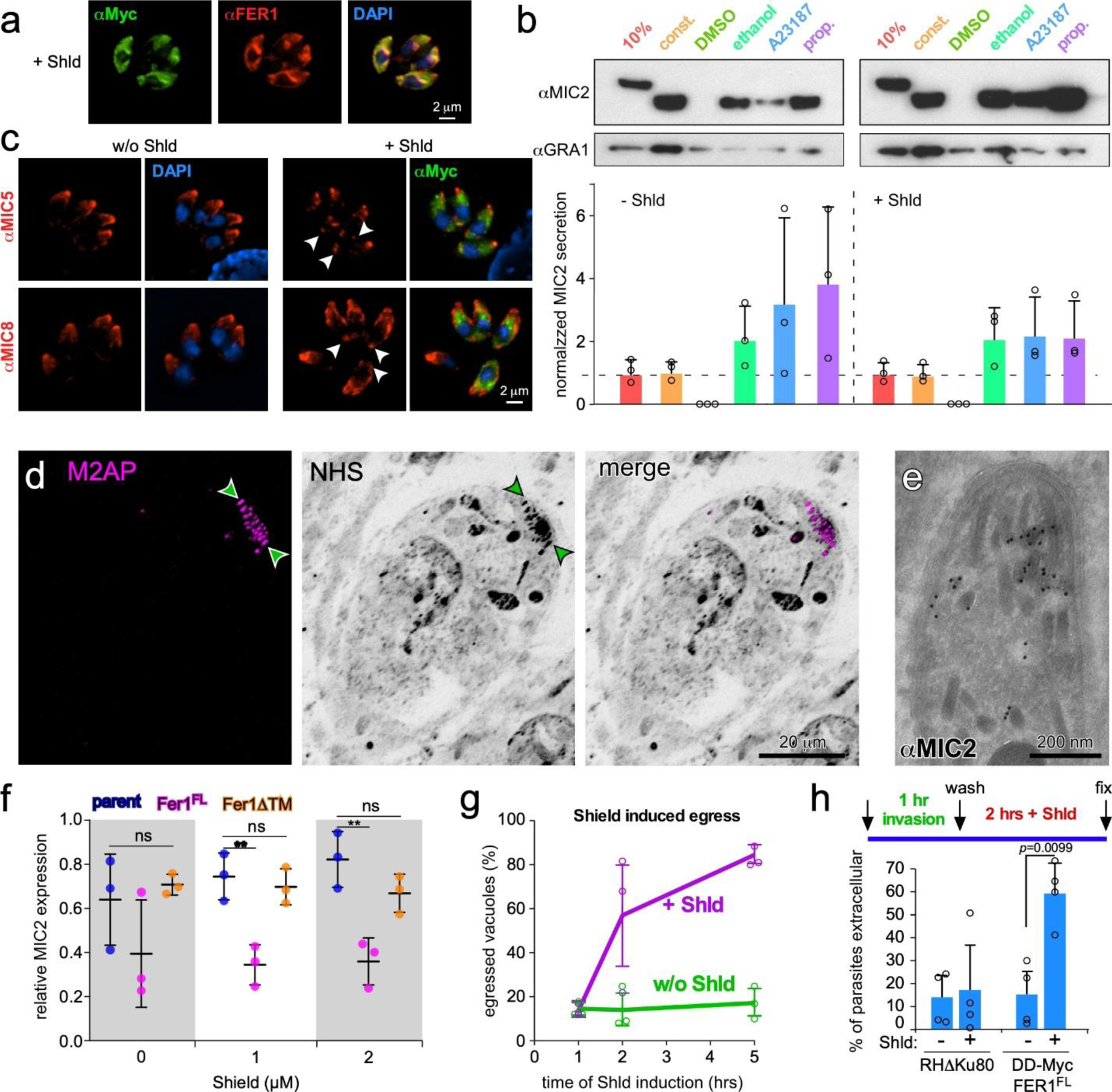
DD-Myc-FER1^FL^ overexpression results in apical microneme migration and a burst of signaling-independent microneme release. (**a**) Co-staining of α-Myc and α-FER1 sera by IFA. Total projection of deconvolved image is shown. (**b**) MIC2 secretion in the supernatant was assessed by western. Parasites were induced 6 hrs with 1 µM Shield-1 and harvested by physical release from the host cell. 10%: 10% of total lysate; const.: 1 hr constitutive secretion at 37°C; 1% ethanol; 2 µM A23187; 500 µM propranolol. DMSO is the vehicle control for A23187. Induced secretion for 5 min at 37°C. Bottom panel represents quantified secretion normalized to the GRA1 signal and to the 10% loading control. n=3+std. Parent line controls in **Suppl Fig S2**. (**c**) Immunofluorescence of DD-Myc-FER1^FL^ co-stained with α-MIC5/α-MIC8 and α-FER1. The Rab5A/C-dependent sub-population becomes scattered upon phenotype induction, although an extreme apical focal point remains. Arrowheads mark microneme signal enriched at the basal end of the parasites. (**d**) Pan-expansion microscopy of DD-YFP-FER1^FL^ parasites induced for 18hrs with 1 µM Shield-1. Compiled stack of 10 slices is shown. Arrowheads indicate the radial set of micronemes. (**e**) IEM of DD-Myc-FER1^FL^ overexpressing parasites stained with MIC2 antibody (10 nm gold particles) induced for 16 hrs with 1 µM Shield-1. (**f**) Quantification of MIC2 protein expression levels assessed by western blot across parent line and mutants using α-tubulin as standard. Quantification was performed using Image Lab software by BioRad. n=3±std. Two-way ANOVA, ** p-value< 0.005. See **Suppl Fig S5** for all western blots. (**g**) Time course of Shield-1 induced egress. Parasites grown for 30 hrs in fibroblasts and induced with 1 µM Shield-1 for the indicated times, fixed with 100% methanol and stained with α-Myc, α-MIC8 and DAPI. Intact vacuoles per field were scored as proxy for egress. n=3±std. (**h**) Modified red-green invasion assay. Top schematic shows parasites were allowed to invade for 1 hr at 37°C followed by a wash to remove non-invaded parasites and 2 hrs 1 µM Shield-1 induction followed by the red-green invasion assay. Lower panel shows the relative number of parasites observed outside the host cell. n=4+std. ANOVA with post-hoc Tukey HSD.

Next, microneme localization and morphology was assessed by IFA, starting with the Rab5A/C-dependent sub-population (MIC5, MIC8). Non-induced parasites display the typical apical microneme distribution. Upon FER1^FL^ overexpression, a modest redistribution of this sub-population to a prominent basal end appeared, while the cortical apical signal became more diffuse (**Fig. 6c**). Staining for the Rab5A/C-independent population (MIC2, M2AP) revealed a dramatically different change in localization pattern. By pan-ExM, the microneme signal was strongly concentrated at the apical end in induced intracellular parasites (**Fig. 6d**; **Fig. 4h**) and correspond to the radial micronemes^12^. Radial micronemes are anchored firmly in the apical region and believed to be primed for exocytosis^11^. To obtain a higher resolution we resorted to both IEM and TEM. Here we observe an accumulation of micronemes at the apical end, confirming the apical MIC signals seen by light microscopy. The densely packed micronemes in the apical region just below the conoid often displayed an elongated morphology (**Fig. 6e**), whereas several micronemes were squeezed inside the conoid (**Suppl Fig. S4**). Further below, we do observe micronemes not stained with MIC2 (**Fig. 6e**), which likely represent the Rab5A/C-dependent sub-population that become scattered as observed by IFA (**Fig. 6c**). Thus, it appears that overexpression of FER1^FL^ has a differential effect on the two microneme sub-populations: the MIC2 population is driven to the apical end, whereas the MIC8 population becomes scattered and accumulates basally. Furthermore, much fewer MIC2 micronemes were observed. We tested relative MIC2 abundance by western blot (**Fig. 6f, Suppl Fig. S5**) in the parental, FER1^FL^ and FER1ΔTM strains and found that FER1^FL^ parasites displayed a significant decrease of MIC2, with negligible changes upon Shield-1 treatment. No changes were observed in FER1ΔTM parasites. These data suggest a role for FER1 in directing microneme sub-populations to different locations, which is a phenomenon not observed before.

To further dissect these phenotypes we tested specific functional capacities. We showed that parasite egress could be triggered by Shield-1 addition in FER1^FL^ overexpressing parasites (**Fig. 6g**). This implies microneme exocytosis as PLP1 released through the micronemes facilitates egress^4^. To determine if FER1 overexpression can trigger microneme exocytosis independently of other signals, we tested if egress could happen in small vacuoles which lacked an acidified PVM^5^ or accumulated secreted diacylglycerol kinase 2 (DGK2) inside the PV^56^, which are both required for egress^6^. To this end we modified the standard red-green invasion assay into an invasion+egress assay exploiting the same differential staining of intracellular vs extracellular parasites^57^. Following 1 hr standard invasion, non-invaded parasites were washed away, and Shield-1 was added for two hrs (**Fig. 6h** top). Nearly 50% of vacuoles with single parasites have egressed (**Fig 6h**. bottom), which largely mimics the 60% egress rate seen after 2 hrs of induction from large vacuoles. Collectively, these data strongly support that Shield-1 induced microneme exocytosis is only due to FER1 overexpression and not due to activation of the signaling pathways leading to egress. This implies that the most apically located radial micronemes act as a readily-releasable vesicle pool.

### 7. Putative Ca^2+^-binding residues are critical for FER1 function

To probe the mechanism of FER1 beyond the TM domain we analyzed the individual C2 domains for their functional potential. C2 domains fold into β-sheets connected by three loops that can bind to proteins or insert in membranes, which can be modulated upon binding Ca^2+^^32,58–60^. Five key residues in the loops, of which the three central amino acids carry most weight^61^, can predict an association with phospholipids or Ca^2+^. Across FER1’s C2 domains, we detected Ca^2+^-binding potential only in the C2D that carries three Asp residues and a supportive Glu in the conserved loop positions to accommodate two Ca^2+^ ions (**Fig. 7a,b**). To perturb the Ca^2+^ binding capacity we mutated all three C2D conserved Asp residues to Ala and conditionally overexpressed this 3DA sequence in the FER1^FL^ and FER1ΔTM DD-YFP constructs (**Fig. 7c,d**). We observed dramatic difference in MIC2 and MIC8 localization across the various mutant alleles, to which end we quantified where in the parasite the signals are distributed (**Fig. 7E-H**). Comparing the MIC2 signal in the overexpressed wild type (wt-3D) with the overexpressed 3DA allele in the FER1^FL^ construct we observed a modest less pronounced apical tip accumulation (**Fig. 7E**). In the ΔTM context, however, the effect of the 3DA mutation is a sharp reduction in overall MIC2 signal that is still somewhat accumulated in the mid-apical region (**Fig. 7G**). For MIC8 in the FER1^FL^ construct, we observe a mild increase in mid apical signal and a depletion of the basal signal in the 3DA allele relative to the wt-3D allele, suggesting MIC8 trafficking into the apical direction (**Fig. 7F**). In the ΔTM construct, we observed a much more variable phenotype for MIC8 in the 3DA allele, which still generally looks like the mid apical accumulation (**Fig. 7H**).

**Figure 7.**
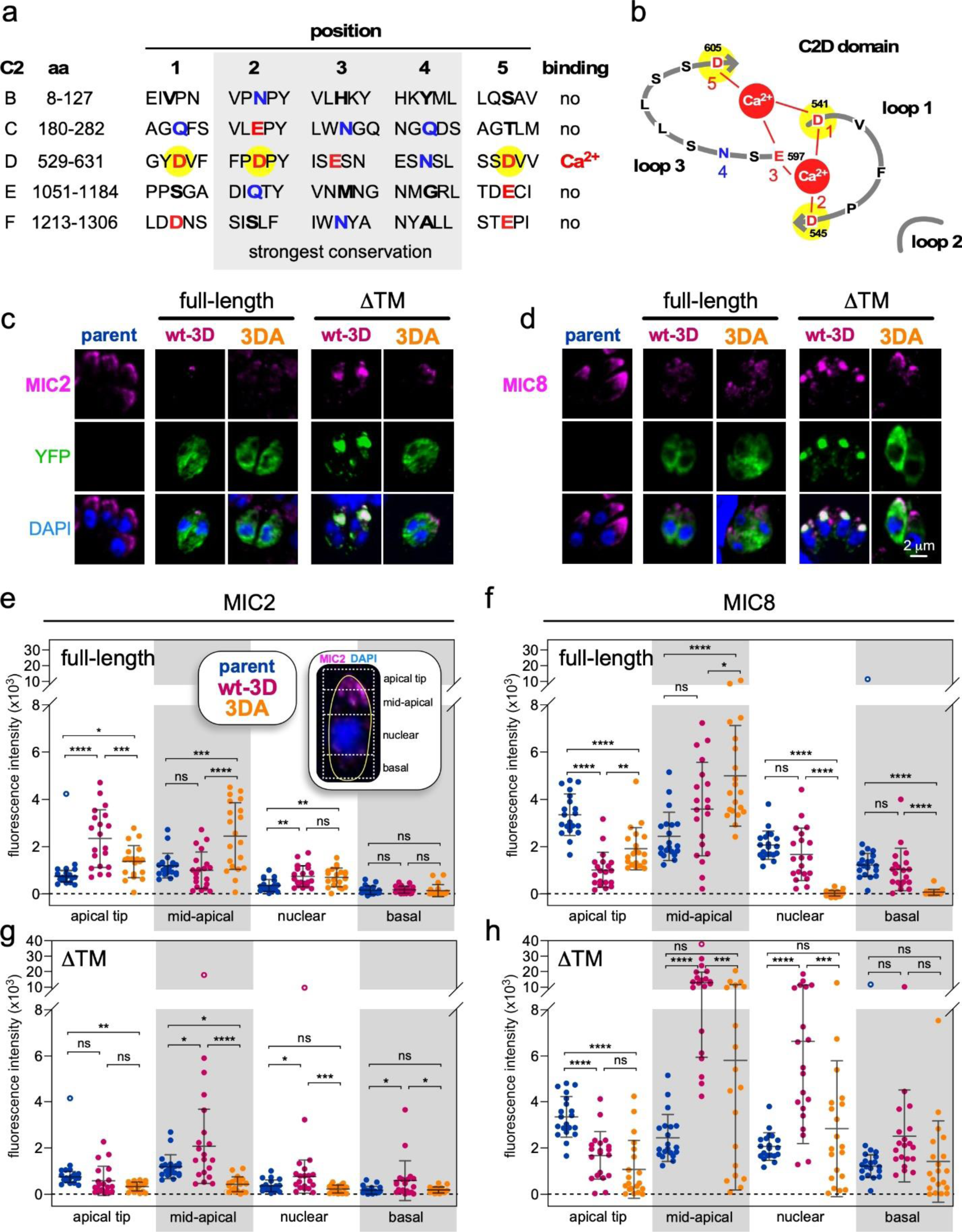
FER1-mediated trafficking of micronemes relies on predicted Ca^2+^-binding residues. (**a**) Sequence analysis of the five conserved key positions (#1-5) in the C2A-F domain loops interfacing with Ca^2+86^. D or E residues (red) stabilize Ca^2+^, N or Q (bleu) are expected to support phospholipid binding. Positions 2, 3, and 4 shaded in grey are more strongly conserved than positions 1 and 5. Yellow highlighted residues were mutated to A to abolish the predicted Ca^2+^-binding capacity in the C2D domain resulting in the mutant FER1(3DA) allele. (**b**) Models of the C2D domain loops and putative Ca^2+^ binding capacity. Yellow highlighted residues as in panel A. (**c, d**) Transient overexpression DD-YFP-FER1ΔTM(3DA) DD-YFP-FER1^FL-^(3DA), respectively, co-stained for YFP, MIC2 and MIC8, as indicated. Each panel contains transfected (YFP positive) and non-transfected (YFP negative) parasite examples. Following electroporation, parasites were allowed to invade for 2 hrs before 1 μM Shield-1 induction for 18 hrs. (**e-h**) Quantification of MIC2 (**e**, **g**) and MIC8 (**f**, **h**) signal localization for wild-type and mutant allele overexpression as indicated. Fluorescence was quantified in the four regions as illustrated in the inserted IFA image in panel E. At least 20 parasites were imaged for each condition. Open symbols represent outlier data points which were excluded from statistical analysis. Error bars display average ±std. One way ANOVA: * *p* < 0.05, ** *p* < 0.01, *** *p* < 0.001, **** *p* < 0.0001. Exact numbers are provided as Data source.

Overall, in FER1^FL^ overexpression the 3DA allele reduces the strong apical shift of MIC2 to the radial micronemes, suggesting that replenishment of the radial micronemes is mediated by the Ca^2+^-binding capacity of FER1. On the other hand, the MIC8 micronemes trend in the other direction away from the very apical end. In the FER1ΔTM overexpressing parasites, MIC2 expression appears to be dependent on Ca^2+^-binding capacity. MIC8 dispersal becomes even more pleomorphic in the 3DA allele and FER1ΔTM context than the wt-3D allele. Thus, the Ca^2+^-binding capacity in the C2D domain mediates apical migration of MIC2 micronemes for putative replenishment and, might regulate MIC2 recycling during division. The MIC8 micronemes, if anything, move in the opposite direction and become dispersed upon disrupting Ca^2+^-binding capacity in the FER1ΔTM context. Collectively, the function of the C2D and TM domains in FER1 confer differential roles on the dynamics of the two microneme populations, both in microneme position and abundance resulting in a very versatile modulating role for FER1 in microneme biology.

## Discussion

The use of a series of FER1 overexpressing DN alleles revealed its multi-faceted role throughout the lytic cycle (**Fig. 8)**.

**Figure 8.**
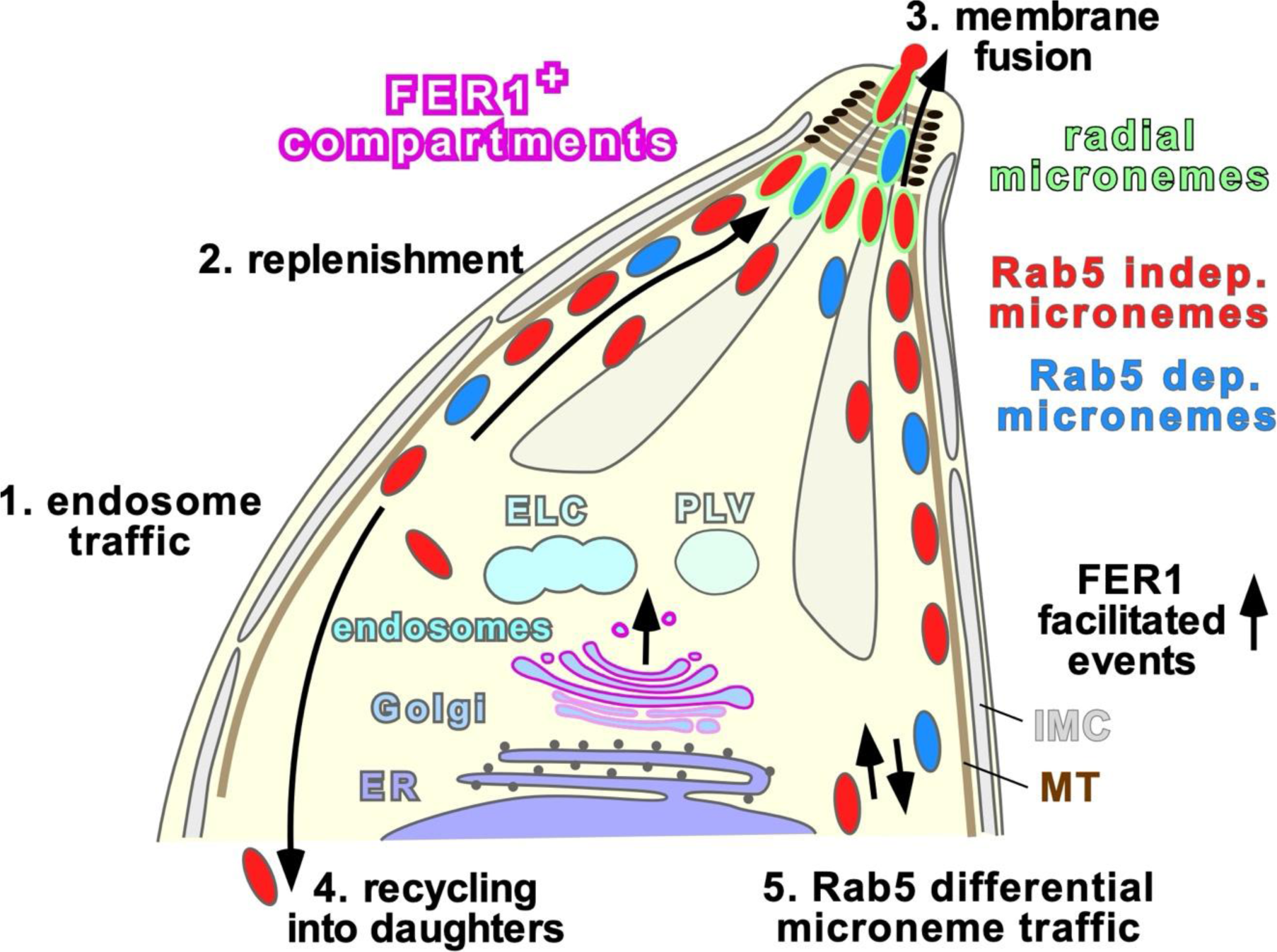
Working model of FER1 mediated microneme events. FER1 mediates the following trafficking steps: 1. from the ELC to the subpellicular cortex (actin dependent?); 2. microneme migration to the apical tip (along the subpellicular microtubules?); 3. microneme membrane fusion with the plasma membrane; 4. recycling of micronemes into the budding daughters (actin dependent); 5. differential trafficking dynamics of the two micronemes populations differentiated by their involvement of Rab5A/C in their biogenesis. We propose that the radial micronemes represent a readily releasable vesicle pool primed for secretion. ELC: endosome-like compartment; IMC: inner membrane complex; MT: microtubules; PLV: plant like vacuole or VAC.

The kinetics of microneme re-direction upon DN-FER1ΔTM overexpression established that matured micronemes are almost instantly released from the cortex, supposedly the subpellicular microtubules, and traffic to a cytoplasmic location. Although similar phenomena were reported upon the disruption of the vacuolar-proton ATPase (v-ATPases) in *Toxoplasma*, they contained immature microneme proteins^25^ unlike what we observed. Also, actin dependent trafficking of micronemes from mother to daughters has been reported^18^, which further supports the feasibility and a biological function for the retrograde microneme organelle transport we observe. Moreover, the effect we see is specific for the micronemes, unlike most other mutants in the endosomal legs of microneme trafficking, which invariable also affect rhoptry protein trafficking^26,29^. Overall, FER1 provides the first mechanistic insight in the microneme trafficking beyond the upstream leg shared with the rhoptries.

Overexpression of FER1^FL^ supports a direct role in membrane fusion in the actual exocytosis. The first piece of evidence is that Shield-1 induction leads to fast parasite egress (**Fig. 6g,h**). In parallel to other secretory systems^62,63^, we believe that only the primed or so-called readily-releasable microneme pool is released in this mutant, but that additional secretion requires renewed priming (e.g. Ca^2+^-dependent events such as potential phosphorylation of FER1^64^). The radial micronemes, accumulated right below the conoid, make a good candidate for this readily-releasable microneme pool^12,13^. Radial micronemes are tightly anchored as they were the only set of micronemes remaining upon VPS9 knock-down^11^. However, we cannot exclude that only one or two micronemes are pre-docked and engaged to complete fusion with the plasma membrane.

MIC2 microneme population squeezes into the apical end of the parasite, indicating anterograde transport along the cortex was triggered as well. A similar microneme apical migration was observed upon knock out of AP1, although that was not unique to the micronemes and acted much more widely across many aspects of vesicular trafficking^29^. Since the AP1 mutant did not lead to microneme exocytosis, we conclude that the secretion burst, and apical movement are independent events facilitated by FER1 overexpression. We interpret the apical movement as part of the process of secretory vesicle replenishment. By comparison, in mammals a role for otoferlin in replenishment of synaptic vesicles is supported, which mimics this particular function of FER1 in *T.* gondii^13,42,65^.

The mis-localizing micronemes upon overexpression of the ΔTM allele, make us assert that the TM domain is associated with microneme organelle trafficking. Further support is provided by the vesicular localization in the gametocyte cytoplasm reported for the FER1 ortholog PbFLP in *P.* berghei^36^. C2 domains appear to directly engage the micronemes. Analysis of FER1 C2 domains pointed at C2D with potential for Ca^2+^ binding and no indicators of phospholipid binding. Among mammals, the C2D domain of otoferlin has been demonstrated to bind Ca^2+39^ and interacts with MyosinVI^66^ allowing vesicle transport^40^. This engagement connects to the actin-myosin dependent microneme recycling during cell division^18^, although there is currently no indication this process is Ca^2+^-dependent. In general, low Ca^2+^ promotes intramolecular protein-protein interactions among otoferlin C2 domains, whereas high Ca^2+^ triggers interaction with phospholipids^40^ and SNARE proteins *in vitro*^67,68^. For *Toxoplasma* FER1, we suggest that the C2 domains function in protein-protein interactions to expose or hide functional domains upon signaling conditions. Indeed, we observe distinct effects on the two microneme sub-populations, indicating that these micronemes have different proteins exposed on their cytoplasmic facing membrane. *Plasmodium falciparum* also harbors two different microneme populations, with different contents that, unlikely *T. gondii*, display distinct exocytosis kinetics^69^. How *P. falciparum* regulates this or whether protein trafficking is differentiated by Rab5A is currently unknown. In future work we will pursue the nature of this microneme specific biology.

In conclusion, FER1 is a key player in several different aspects of microneme biology. FER1’s Golgi-endosomal localization suggests it is involved in microneme biogenesis, likely at the vesicular trafficking level. In addition, it can direct microneme organelle traffic differentially based on microneme population as well as differentially during microneme secretion versus cell division. Finally, FER1 has a direct role in membrane fusion facilitating exocytosis. Thus, FER1 is a versatile player that may be a tool to understand the intricate details of microneme biology.

## Material and Methods

### Parasites and host cells

Transgenic derivatives of the RH strain were maintained in human foreskin fibroblasts (HFF) or hTERT immortalized HFF cells as previously described^70^. Parasite transfections and selections use 1 μM pyrimethamine, 20 µM chloramphenicol, 5 µg/ml FUDR, or a combination of 25 mg/ml mycophenolic acid and 50 mg/ml xanthine (MPA/X). All parasite lines were cloned by limiting dilution.

### Generation of constructs and parasite lines

All primer sequences are provided in **Supplementary Table S1**; all plasmids used are provided in **Supplementary Table S2**. Expression plasmids fusing ddFKBP destabilization domain (DD) with FER1 were generated from tub-DD-Myc-YFP/sagCAT plasmid^52^ by replacing YFP with the PCR amplified FER1 CDS (primer pair 1573/1574) by *Avr*II and *Xma*I restriction enzymes to generate tub-DD-Myc-FER1^FL^/sagCAT and in tub DD-YFP-TgNek1-2(MCS)/sagCAT^71^ to generate tub-DD-YFP-FER1^FL^/sagCAT. FER1ΔTM constructs were generated by amplifying a 3’ section without TM domain (primer pair 1651/1652; deletion of the C-terminal 21 aa) and cloning the product in the FER1-FL plasmids using *Nhe*I and *Xma*I. The FER1 Ca^2+^-binding mutants in the C2D domain were generated by Q5 site directed mutagenesis kit (NEB) using primers 4833/4834 to change positions A1622 and A1634 to C resulting in Asp codon 542, 545 changes to Ala, and primers 5892/5893 to change position A1814 to C resulting in Asp codon 605 to Ala.

Plasmid tub-YFP-FER1(TM) encoding only the 31 most C-terminal aa of FER1 including the TM domain was cloned by PCR amplification using primer pair 4786/4788 and cloned by Gibson assembly into *Bgl*II/*Avr*II digested tub-YFPYFP/sagCAT plasmid^34^.

Plasmid tub-IMC3mCherry/DHFR was cloned by swapping IMC3mCherry from tub-IMC3mCherry/sagCAT^72^ with *Pme*I/*Avr*II into tub-YFPYFP(MCS)/DHFR^34^.

To generate pmic3-MIC3-Cherry/DHFR 1.3 kb of promoter region together with 1.2 kb of the ORF encoding genomic locus was PCR amplified from genomic DNA (primer pair 4864/4865) and Gibson assembled into *Pme*I and *Avr*II digested tub-mCherry_2_/DHFR^73^.

Parasites expressing endogenously tagged SORTLR_Ollas/HXGPRT were generated by inserting a PCR fragment (amplified with primer pair 6014/6015) targeting the 3’-SORTLR locus using a SmOllas/HXGPRT template. Insertion at the C-terminus was facilitated by CRISPR/Cas9 using a pU6 plasmid^74^ containing primer 6013 as guide RNA.The SmOllas/HXGPRT template was produced by amplifying the SmOLLAS from the pCAG_mRuby2_smFP OLLAS plasmid (Addgene cat no. 59761^75^) using primer pair 5688/5689, while the HXGPRT resistance cassette was amplified from the mAID-3xMyc-3’hxgprt-5’dhfr-HXGPRT-3’dhfr plasmid^76^ (primer pair 5690/4769). PCR fragments were cloned into the previously *NotI* and *KpnI* digested ptub-YFP-YFP/sagCAT plasmid^72,77^, via Gibson assembly (NEBuilder® HiFi 478 DNA Assembly Master Mix; NEB) (**Suppl Fig. S7**).

### Antiserum generation

TgFER1 amino acids 669-877 including the diverse C2 domain DE were PCR amplified using primers Ava-LIC-Fer1-F/R and fused to a 6xHis tag in plasmid pAVA0421^78^, expressed in *Escherichia coli* BL21, purified by Ni-NTA chromatography (Invitrogen), and used to immunize a guinea pig (Covance, Inc). Serum was affinity purified as described previously^79^ against recombinant His6-TgFER1.

### Live-cell microscopy

For monitoring egress (P30-YFP and GCaMP3 expression), parasites were grown in hTERT confluent 15 mm glass bottom cell culture dish (MatTek Corporation, cat #801002) for 30 hrs and then induced with 2 µM Shield-1 for 90 min at 37°C. Dishes were live cell imaged on a Zeiss Axiovert 200M inverted microscope for 15 min at 2 images per minute. To monitor DD-YFP-FER1ΔTM accumulation a Leica TCS SP5 scanning confocal microscope with incubation chamber in the Boston College Imaging Core in consultation with Bret Judson. Upon addition of 1 µM Shield-1 images were captured every 5 mins for 3 hrs. All images were acquired, analyzed and adjusted using Leica, Volocity (Quorum Technologies) and/or ImageJ/FIJI software^80,81^.

### (Immuno-) fluorescence microscopy

Indirect immunofluorescence assays were performed on intracellular parasites grown for 18 hrs in 6-well plate containing coverslips confluent with HFF cells fixed with 100% methanol (unless stated otherwise) using primary antisera as listed in **Supplementary Table S3**. Alexa 488 (A488) or A594 conjugated goat α-mouse, α-rabbit, α-rat, or α-guinea pig were used as secondary antibodies (1:500, Invitrogen). DNA was stained with 4’,6-diamidino-2-phenylindole (DAPI).

For intracellular IFAs, parasites were allowed to invade and replicate for 24 h after which 1 µM Shield-1 was applied for 18 hr (DD-Myc/YFP-FERΔTM) or 2 µM Shield-1 for 3 hr (DD-Myc-FER^FL^). Extracellular parasites grown ± Shield-1 for 18 hr were harvested by mechanical lysis and captured on Poly-L-lysine coated coverslips. A Zeiss Axiovert 200 M wide-field fluorescence microscope was used to collect images, which were deconvolved and adjusted for phase contrast using Volocity software). Co localization assays were imaged on a Zeiss AxioImager Z2 microscope with a sCMOS camera (Flash 4, Hamamatsu) using an Apotome2. Apotome images were processed in ZEN Blue using automatic processing while Pearson Correlation Coefficient (PCC) were measure using ImageJ.

### Pan expansion microscopy

Pan expansion microscopy assays were performed basically as described^82,83^. Intracellular parasites were grown for 18 hrs in 6-well plate containing HFF cells confluent coverslips. and fixed with 4% PFA for 20 min at RT. Samples were then incubated in the 2× fixing/anchoring solution (2% acrylamide; 1.4% formaldehyde in 1×PBS) for 5 h at 37 °C in a dessicator box. Excess fixing/anchoring solution was washed with 1xPBS before sealing the coverslips in the gelation chamber (as described in^82^). 90 µl of first gelation solution (19% sodium acrylate; 10% acrylamide; 0.1% DHEBA; 0.25% TEMED; 0.25% APS in 1xPBS) was added directly on the cells, before covering with a new coverslip that becomes the lid inducing the formation of a homogeneous height gel. The gelation chamber was incubated 15 min at RT, then placed in a humid chamber for 1.5 hrs at 37°C. After gelation, the coverslips were removed from the gelation chamber and placed in a 6-well plate filled with 2 ml of denaturation buffer (200 mM SDS; 200 mM NaCl; 50 mM Tris; pH 6.8) and incubated for 15 min at RT while shaking until the gel detached from the coverslip. Next, gels were transferred in a 2 ml Eppendorf tube with 1 ml fresh denaturation buffer and incubated for 1 hr at 37°C. After denaturation, the gels were placed in a Petri dish, washed once with 1xPBS before starting the first expansion with ddH_2_O. Water was replaced three times in 30 min intervals before full expansion overnight. Gels were cut in approximately 1 cm² pieces and placed in a 6-well plate filled with 1.5 ml neutral hydrogel solution (10% acrylamide; 0.05% DHEBA; 0.05% TEMED; 0.05% APS in ddH_2_O) and incubated 2×20 min at RT while shaking. Excess monomer solution was removed by pressing carefully with Kimwipes on both sides of the gel. Gels were placed on a microscope slide and covered with a coverslip before incubation in a humid chamber for 1 hr at 37°C.

For the second expansion, gels were placed in a 6-well plate and washed once with 1xPBS before two incubation with 1 ml second gelation solution (19% sodium acrylate; 10% acrylamide; 0.1% bis-acrylamide; 0.05% TEMED; 0.05% APS in 1xPBS) for 15 min rocking on ice. Gels were placed on slides and excess solution removed as described above, then covered with coverslips and places in humid chamber for 1 hr at 37°C. Finally, the first and neutral gels were dissolved by 2 ml of 0.2 M NaOH for 1 hr at RT while shaking. Dissolution was followed by several washes with 1xPBS until pH becomes neutral. For antibody staining, small pieces of gel were cut and incubated with α-M2AP (1:250) followed by AlexaFluor 594 (1:200) antibodies, overnight at RT followed by 3 hrs at 37°C the next day. Antibodies were diluted in 2% BSA and washed 3×20 min with PBS-Tween (0.1%). Finally, NHS-ester 488 staining was performed at 20 µg/ml in 1xPBS for 1 hr. Washing was performed as described above, before a final expansion overnight in ddH_2_0. Gels were imaged in 35 mm Mattek on a Zeiss LSM 880 using Airyscan and a Plan-APOCHROMAT (x40; water). Z-stacks were collected for all gels using 0.2 µm step size through the entire depth of parasites. Post processing of Airyscan images was completed in ZEN Blue and ImageJ.

### Electron microscopy

For ultrastructural observations of *T. gondii*-infected cells by thin-section transmission electron microscopy (TEM), infected cells were fixed in 2.5% glutaraldehyde in 0.1 mM sodium cacodylate (EMS) and processed as described^84^. Ultrathin sections of infected cells were stained before examination with a Philips CM120 EM (Eindhoven, the Netherlands) under 80 kV. For immunoelectron microscopy (IEM) samples were prepared as described before^34^. Sections were immunolabeled with MIC2 MAb 6D10 in 1% fish skin gelatin and then with goat anti-IgG antibodies, followed by 10-nm protein A-gold particles before examination with a Philips CM120 electron microscope under 80 kV.

### Host cell invasion

Extracellular parasites from 80% naturally lysed flask were induced with 2 µM Shield-1 for 90 min at 37°C before they were allowed to invade HFF host cells for 10 min at 37°C^7^. The red/green invasion assay was performed as described^85^ using Alexa594- and Alexa488-conjugated SAG1 MAb T41E5. Three images were taken per biological replicate (3 biological replicates total) on an EVOS FL (Life Technologies). The number of invaded versus uninvaded parasites were enumerated manually for at least 300 parasites per counted sample.

### Secretagogue induced egress

The egress assay was performed essentially as described previously^34^. Parasites were grown in HFF monolayers for 30 h after which the phenotype was induced with 1 µM Shield-1 for 18 hrs. Egress was triggered with 1-3 µM A23187, 500 µM propranolol, 50-150 µM BIPPO (kindly shared by Dr. Jeff Dvorin) 0.01-0.1% saponin or DMSO for 5 min at 37°C, followed by IFA with IMC3 and GRA3 antisera. Egressed, permeabilized and intact vacuoles were counted^56^. 100 vacuoles were counted for each experiment and three biological replicates were performed.

### Shield-1 induced egress

Parasites were inoculated on HFF coverslips and allowed to grow for 30 hrs and then induced with either 1 µM Shield-1 for 2 hrs or 1 µM calcium ionophore A23187 for 5 min as a control, prior to fixation with 100% methanol and IFA staining using MAb 9E10 cMyc and rabbit αMIC8 (antisera details in **Supplementary Table S1**). DNA was stained with 4’,6-diamidino-2-phenylindole (DAPI). The number of intact vacuoles per 20 fields was enumerated. Three independent biological replicates per experiment were performed.

### Combined invasion-egress Assay

Parasites were grown for 36 hours, mechanically lysed in standard ED1 parasite medium and allowed to invade coverslips coated in an HFF monolayer for 1 hr. All unattached parasites were then washed off with a PBS and coverslips were incubated in 1 μM Shield-1 or vehicle control for an additional 2 hrs. Coverslips were then fixed with PFA and IFA was performed as described for a typical red-green invasion assay. The number of parasites was counted in 10 fields and four independent biological replicates per experiment were performed.

### Microneme secretion by Western blotting

Microneme secretion by western blot was performed as published^16^. Freshly lysed parasites were resuspended in DMEM/FBS and transferred to a 96-well polystyrene round-bottom plate (CELLTREAT Scientific Products). Secretion was induced by 1-3 µM A23187, 500 µM propranolol, 1% ethanol or DMSO for 5 min at 37°C. Constitutive microneme secretion was assessed by incubation without secretagogue at 37°C for 60 min. Supernatants were probed by western blot using 6D10 MIC2 MAb and TG17.43 GRA1 MAb and HRP conjugated secondary antiserum. Signals were quantified using a densitometer. Three independent biological replicates per experiment were performed.

### Microneme abundance by Western blotting

Intracellular parasites were stimulated overnight with 0, 1 or 2 µM Shield-1 (AOBIOUS) at 37°C. The day after, intracellular parasites were washed with DMEM/FBS and released from host cell using a 26G needle. Parasites were resuspended in RIPA buffer and probed by western blot using 6D10 MIC2 Mab and 12G10 Tubulin Mab and HRP conjugated secondary antiserum. Signals were quantified for three independent biological replicates.

### Microneme secretion by IFA

IFA on parasites exposed to a host cell monolayer was performed as reported^34^. Parasites were resuspended in Endo buffer and spun onto HFF cells in a 6-well plate at 28*g, 5 min, RT and allowed to settle for 20 min at 37°C. Secretion was induced by replacing the buffer with 3% FBS in DMEM and 10 mM HEPES (pH 7.2) and incubation at 37°C for 5 min. PBS-washed coverslips were fixed with 4% formaldehyde and 0.02% glutaraldehyde, and subjected to IFA with anti-MIC3, -MIC5, -MIC8 or MIC10 in the presence of 0.02% saponin.

### Gliding motility trail assay

Trail assays were performed as previously described^28^. Parasites were induced with 1 µM Shield-1 for 18 hr, mechanically released, resuspended in ED1 with 1 µM Shield-1 and incubated on poly-L-lysine coated coverslips for 15 min at 37°C. Parasites were fixed with 4% formaldehyde and 0.02% glutaraldehyde and stained with DG52 SAG1 MAb (64) to visualize trails.

### Statistics

Student’s paired *t*-test, one-way and two-way analysis of variance (ANOVA) using post hoc Tukey correction were performed.

## Acknowledgements

We thank Bret Judson and Dr. Patrick Autissier of the Boston College Imaging and Flow Cytometry Cores, respectively, for infrastructure and support, Dr. Sander Groffen for assistance with molecular modeling, Klemens Engelberg for establishing the *Toxoplasma* pan-ExM protocol, Emily Stoneburner, Natalie Sandlin, and Elizabeth Gray for technical support, Drs. Gustavo Arrizabalaga, Peter Bradley, Vern Carruthers, Iain Cheeseman, Jean-François Dubremetz, Wassim Daher, Jeff Dvorin, Maryse Lebrun, Sebastian Lourido, Sabrina Marion, Silvia Moreno, Naomi Morrissette, Jeroen Saeij, David Sibley, and Gary Ward for sharing reagents. We also thank Kim Zichichi at the EM Facility at Yale University for processing and immunolabeling our samples.

This study was supported by National Science Foundation (NSF) Major Research Instrumentation grant 1626072, National Institutes of Health grants AI108251 (B.I.C.), AI060767 (I.C.), AI122042 (M.-J.G), AI099658 (M.-J.G.), and AI122923 (M.-J.G.), and American Cancer Society grant RSG-12-175-01-MPC (M.-J.G.).

## Author contributions

DNAT performed all functional experiments on the DD-[Myc/YFP]-FER1ΔTM parasites, imaging of the DD-Myc-FER1^FL^ line, and time lapse microscopy, ARES performed αFER1 localizations, pan-ExM, MIC2 quantification by western blot, generated DDD-AAA mutants and quantified their phenotypes, AAD performed all functional studies on the DD-Myc-FER1^FL^ parasites, IC performed all electron microscopy studies, BIC cloned FER1 cDNA, established the overexpression plasmids and assessed the initial phenotype, DNAT, ARES, AAD and MJG designed the experiments and interpreted the data, ARES and MJG wrote the manuscript and all authors reviewed and edited the manuscript.

## Supplementary information

**Supplementary Movie S1.** Time lapse experiment of DD-YFP-FER1ΔTM parasites co-transfected with MIC3-mCherry to track the micronemes. At t=0, 2 μM Shield-1 was added and images collected every 5 minutes. Scale bars 80 μm.

**Supplementary movie S2.** Time lapse experiment of DD-YFP-FER1ΔTM parasites co-transfected with IMC3-mCherry to track cell division status. At t=0, 2 μM Shield-1 was added and images collected every 5 minutes. Scale bars 80 μm.

---

**Supplementary Figure S1.**
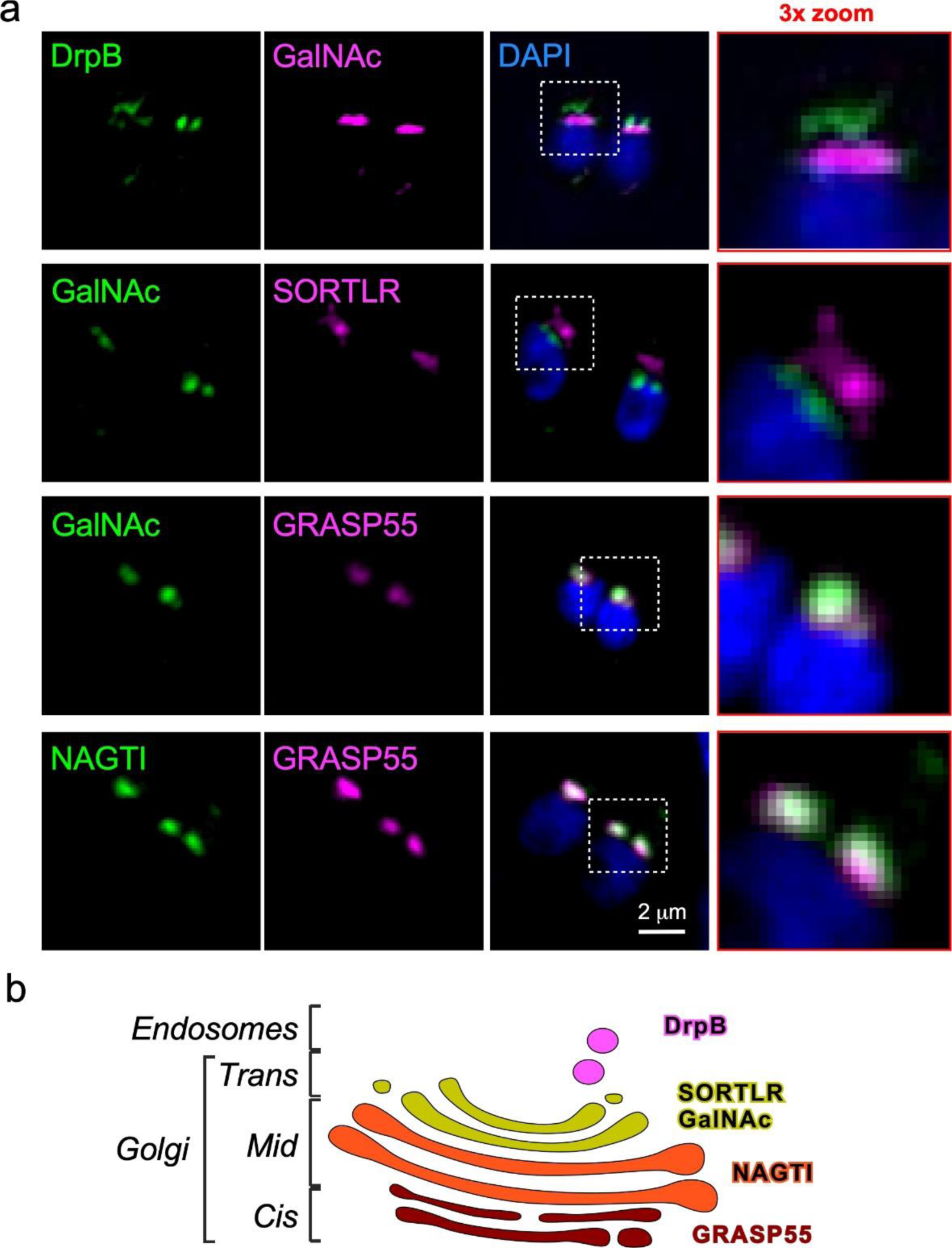
Calibration of Golgi and endosome markers. (**a**) Co-transfected parasites with the markers as indicated. (**b**) Schematic representation of collected insights.

**Supplementary Figure S2.**
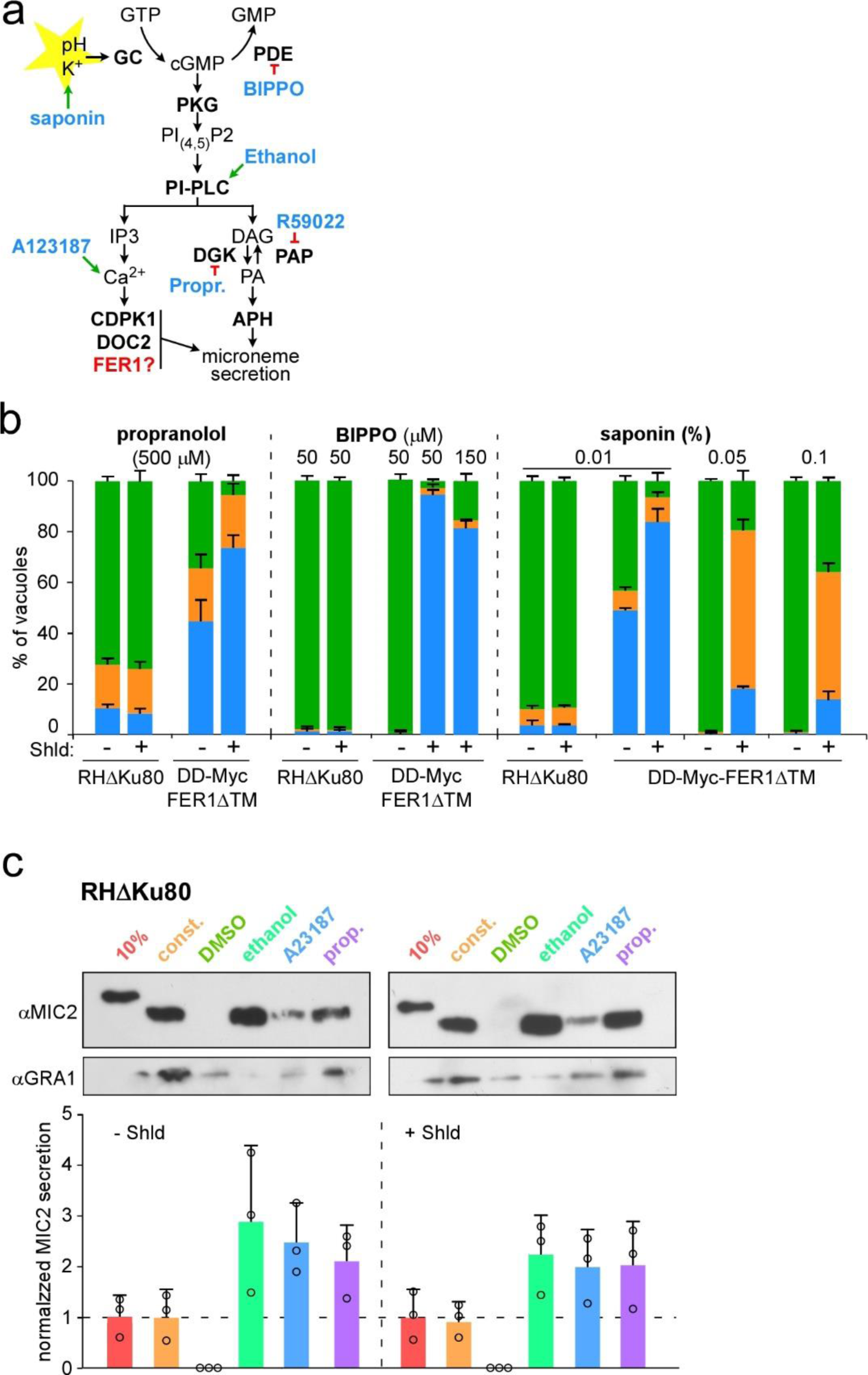
Egress assays and microneme secretion. (**a**) Schematic overview of signaling toward microneme secretion and egress, highlighting pharmacological agents acting on various events. (**b**) Induced egress assays. Parasites grown for 30 hrs in fibroblasts and induced for 18 hrs with 1 µM Shield-1 were triggered for egress with the pharmacologicals as indicated, fixed and stained with α-IMC3 (parasite cortex) and α-GRA3 (PVM) sera and scored for status of vacuole permeabilization and/or egress. n=3±std. Parasites were not under Shield-1 pressure during the 5 min pharmacological incubation. (**c**) Assessment of microneme secretion by western blotting by detecting MIC2 release in the supernatant under various triggers. Parasites were induced 18 hrs with 1 µM Shield-1 and harvested by physical release from the host cell. 10%: 10% of total lysate; const.: 1 hr constitutive secretion at 37°C in absence of secretagogue; 1% ethanol; 2 µM A23187; 500 µM propranolol. DMSO is the vehicle control for A23187. Induced secretion for 5 min at 37°C. Bottom of panel represents quantified secretion normalized to the GRA1 signal and to the 10% loading control for each condition. n=3±std.

**Supplementary Figure S3.**
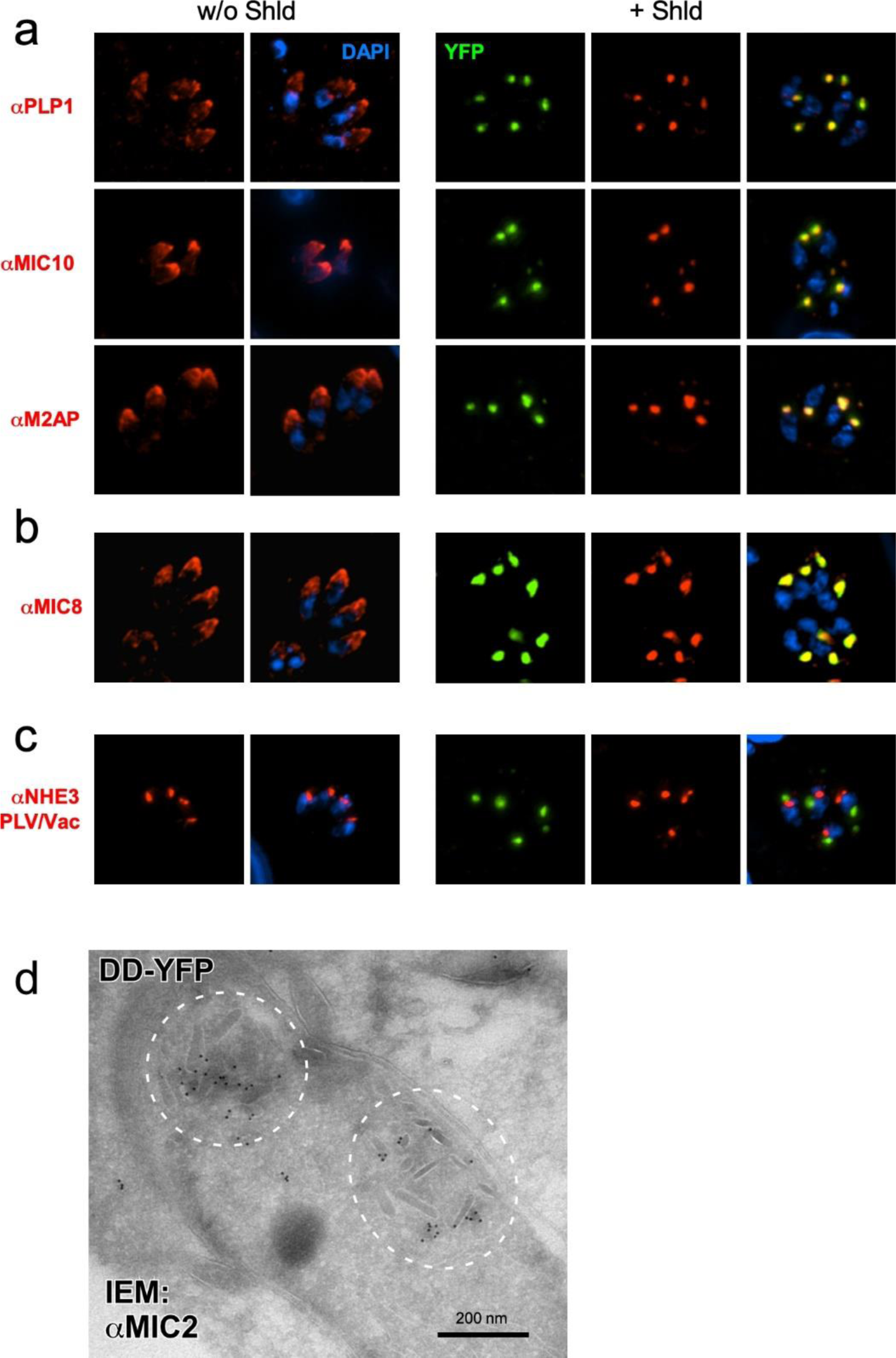
Additional localization data in the DD-YFP-FER1ΔTM dominant negative mutant. (**a**) Additional microneme proteins in the MIC2 group. (**b**) Additional protein in the MIC3/5/8 group. (**c**) NHE3 is a specific marker of the PLV compartment^1^. Parasites were treated with 1 µM Shield-1 for 18 hrs. (**d**) IEM with MIC2 antibody (10 nm gold particles) of DD-YFP-FER1ΔTM overexpressing parasites induced for 7 hrs with 3 µM Shield-1. Dotted circles mark atypical microneme accumulations in the cytosol.

**Supplementary Figure S4.**
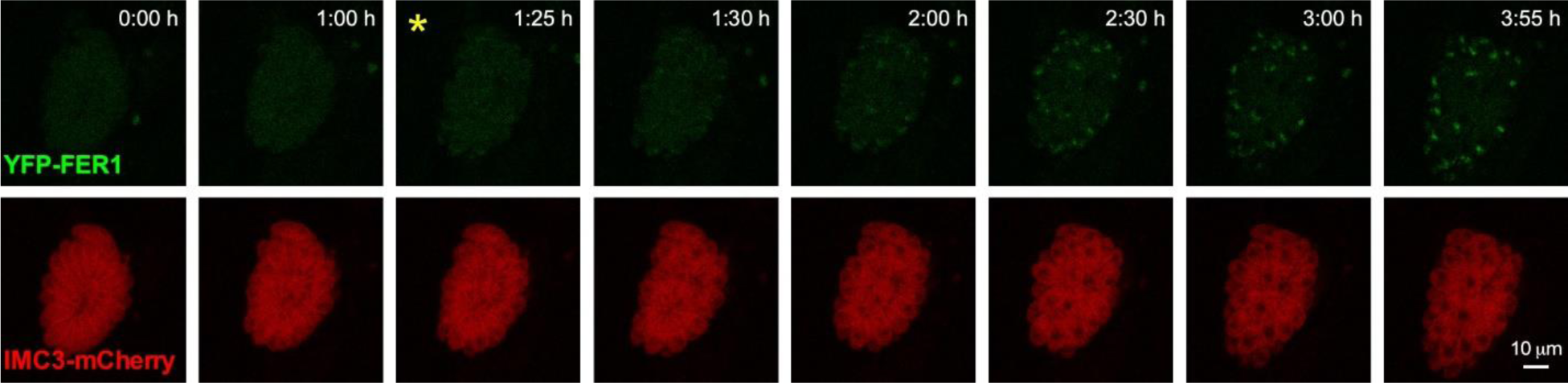
Select panels from time lapse experiment of DD-YFP-FER1ΔTM parasites co-transfected with IMC3-mCherry to track cell division. At t=0, 2 μM Shield-1 was added. The first time point at which the piling up of YFP could be convincingly observed is marked with a yellow asterisk. Panels from supplementary movie S2.

**Supplementary Figure S5.**
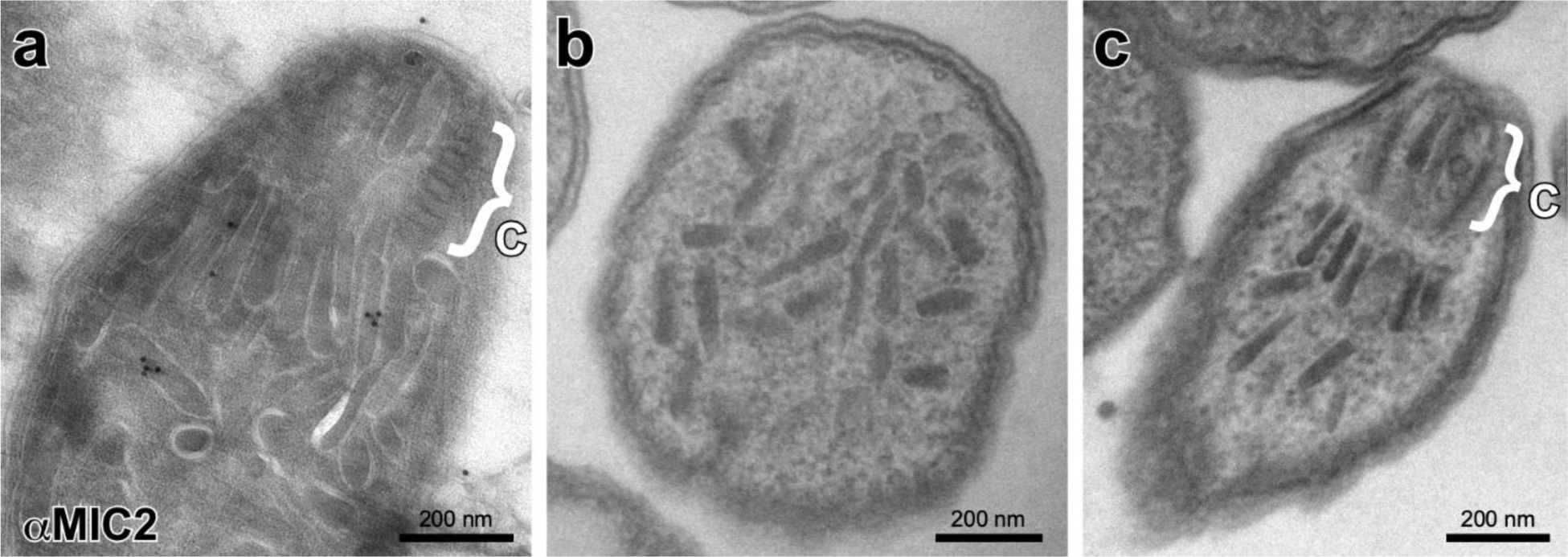
(**a**) IEM of DD-Myc-FER1^FL^ overexpressing parasites stained with MIC2 antibody (10 nm gold particles) induced for 16 hrs with 1 µM Shield-1. MIC signal localizes to apically accumulated micronemes which display a stretched or extended morphology. (**b, c**) TEMs of extracellular DD-Myc-FER1^FL^ overexpressing parasites induced for 3 hrs with 2 µM Shield-1. Cross section in **b** shows the extended feature or the densely packed microneme organelles, whereas **c** shows the radial micronemes just below the conoid, again with a slightly stretched or extended appearance. Accolade brackets labeled ‘C’ mark the conoid.

**Supplementary Figure S6.**
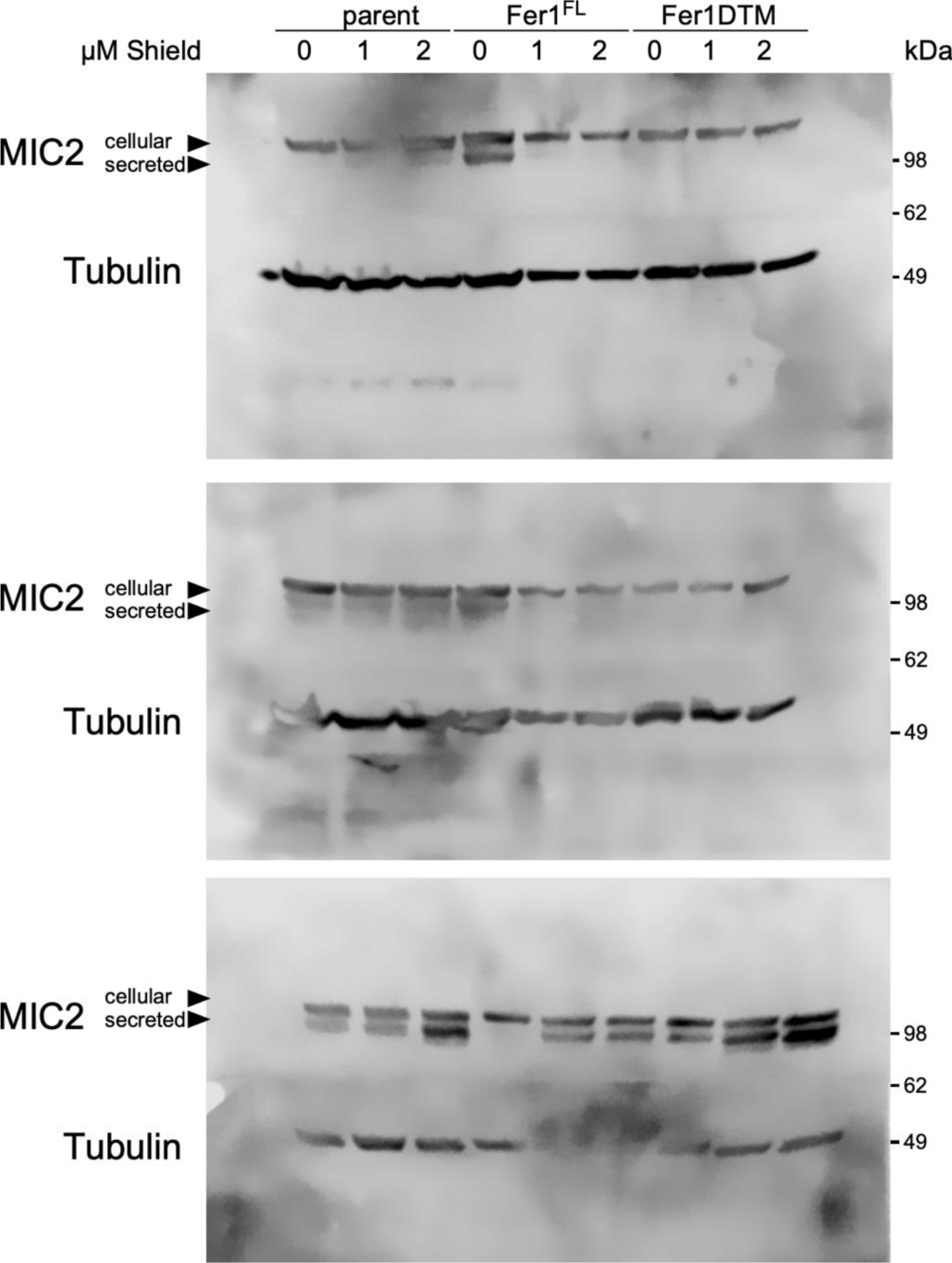
Western blot used to quantify the amount of MIC2 in each parasite strain and under the conditions as indicated. Blots were probed simultaneously with MIC2 and α-tubulin (loading control) antisera.

**Supplementary Figure S7.**
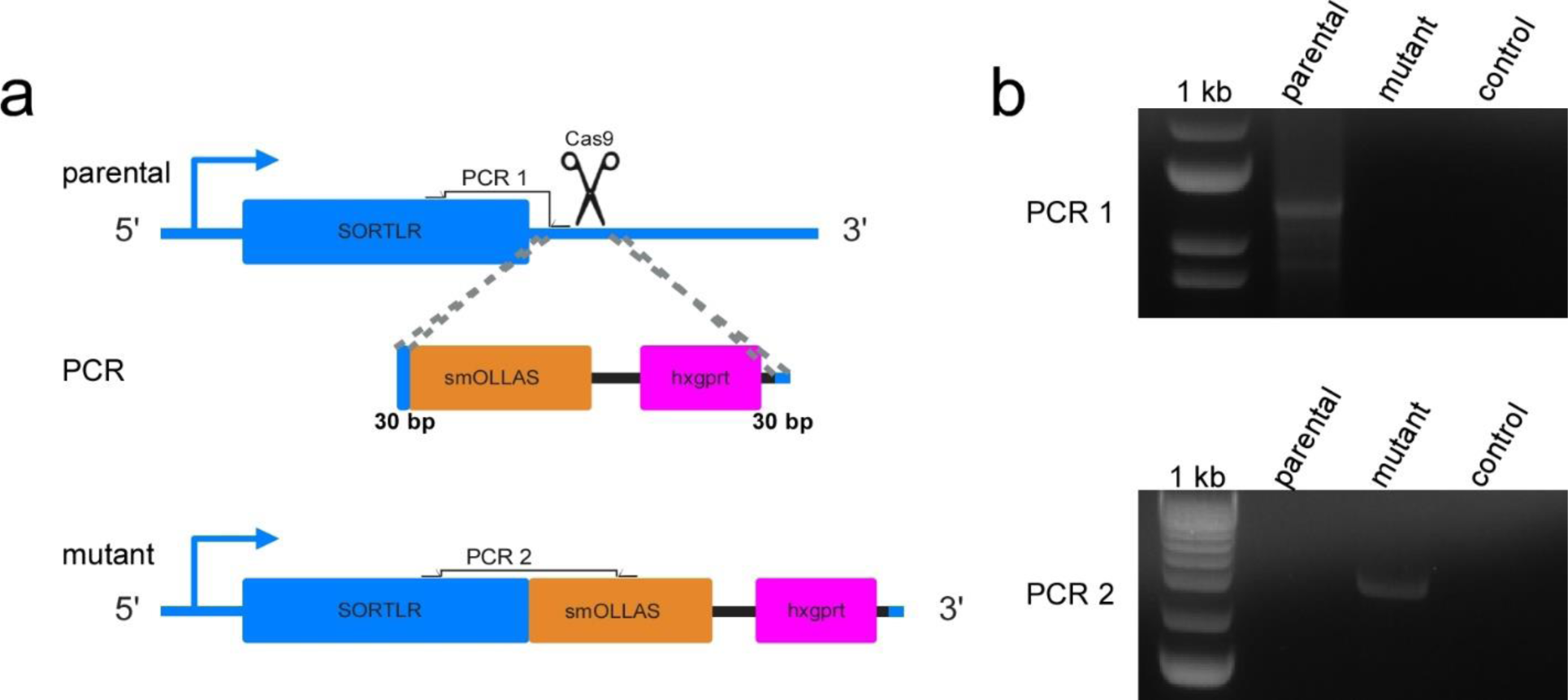
Sortilin_SmOLLAS/RHΔku80 generation and PCR validation. (**a**) Schematic representation of the one step strategy that simultaneously tags sortilin at the C terminus with SmOLLAS tag and introduces the hxgprt selectable marker. Double homologous recombination is facilitated by 30 bp homologous flanks (highlighted in bold) and a CRISPR/Cas9 generated dsDNA break. (**b**) Diagnostic PCR reactions as indicated on parental (RHΔKu80), and clonal strain (mutant), and a no DNA negative control (control). Primer pair localizations are marked in panel a using the following primer pairs: PCR 1: primers 6068 and 6069; PCR 2: primers 6068 and 6070; (see **Supplementary Table S1** for primer sequences). Expected lengths are respectively: 1225 and 2371 bp.

**Supplementary Table S1.**
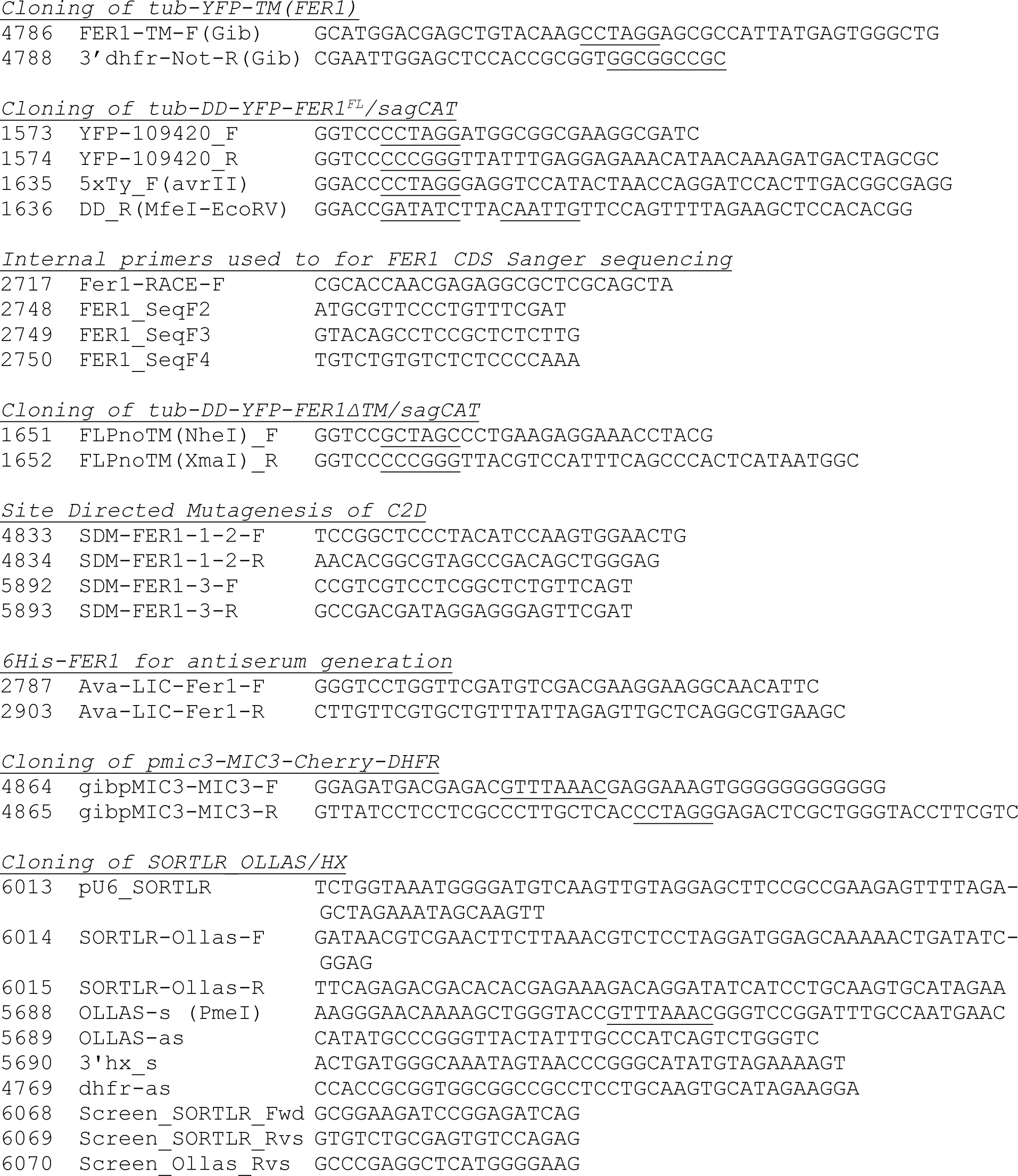
Description of oligonucleotides used. Restriction enzyme sites underlined.

**Supplementary Table S2.** Description of plasmids used.

| Plasmid | Source and reference |
| --- | --- |
| mAID-3xMyc-3'hxgprt-5'dhfr-HXGPRT-3'dhfr | in house<sup>2</sup> |
| pCAG_mRuby2_smFP OLLAS | Addgene<sup>3</sup> |
| ptub-DD-YFP-FER1ΔTM/sagCAT | this paper |
| ptub-DD-Myc-FER1ΔTM/sagCAT | this paper |
| ptub-DD-Myc-FER1ΔTM(<sup>3DA</sup>)/sagCAT | this paper |
| ptub-DD-Myc-FER1<sup>FL</sup>/sagCAT | this paper |
| ptub-DD-Myc-FER1<sup>FL</sup>(<sup>3DA</sup>)/sagCAT | this paper |
| ptub-YFP-YFP(MCS)/sagCAT | in house<sup>4</sup> |
| ptub-YFP-TM<sup>FER1</sup>/sagCAT | this paper |
| ptub-IMC3-mCherryRFP/sagCAT | in house<sup>5</sup> |
| pmic3-MIC3-mCherryRFP/hxgprtDHFR | this paper |
| ptub-GalNAc-YFP/sagCAT | David S. Roos<sup>6</sup> |
| ptub-GRASP55-RFP/sagCAT | David S. Roos<sup>7</sup> |
| ptub-NAGTI-YFP/sagCAT | David S. Roos<sup>6,7</sup> |
| Rab5A-HA OE Transient overexpression | Sabrina Marion<sup>8</sup> |
| Rab7-HA OE Transient overexpression | Sabrina Marion<sup>8</sup> |

**Supplementary Table S3.** Description of antibody and antisera used.

| **Name**<br>**(monoclonal)** | **Species** | **Dilution**<br>**(IFA)** | **Dilution**<br>**(western)** | **Source and reference** |
| --- | --- | --- | --- | --- |
| hCentrin2 | rabbit | 1:1000 |  | Iain Cheeseman, unpublished |
| DrpB | rat | 1:2000 |  | Peter Bradley<sup>9</sup> |
| HA (3F10) | rat | 1:3000 |  | Roche |
| GFP | rabbit | 1:500 |  | Torrey Pines Biolabs |
| GRA1 (TG17.43) | mouse | 1:500 | 1:15000 | BioVision.com<sup>10</sup> |
| GRA3 (T62ch) | mouse | 1:500 |  | Maryse Lebrun, Jean-François Dubremetz<sup>11</sup> |
| FER1 | guinea pig | 1:1000 | 1:1000 | This paper |
| IMC3 | rat | 1:2000 | 1:1000 | In house<sup>5</sup> |
| cMyc (9E10) | mouse | 1:50 | 1:50 | Santa Cruz Biotech |
| MIC2 (6D10) | mouse | 1:2000 | 1:8000 | David Sibley<sup>12</sup> |
| MIC2 | rabbit | 1:1000 | 1:8000 | David Sibley<sup>12</sup> |
| MIC3 | rabbit | 1:100 | 1:400 | Jean-François Dubremetz<sup>13</sup> |
| MIC5 | rabbit | 1:500 | 1:10000 | Vern Carruthers<sup>14</sup> |
| MIC8 | rabbit | 1:1000 |  | Dominique Soldati<sup>15</sup> |
| MIC10 | rabbit | 1:500 | 1:15000 | Vern Carruthers<sup>16</sup> |
| M2AP | rabbit | 1:1000 | 1:1000 | Vern Carruthers<sup>17</sup> |
| NHE3 | guinea pig | 1:1500 |  | Gustavo Arrizabalaga<sup>1</sup> |
| OLLAS | rat | 1:1000 |  | Invitrogen |
| proM2AP | rabbit | 1:1000 | 1:1000 | Vern Carruthers<sup>17</sup> |
| proMIC5 | rabbit | 1:500 |  | Vern Carruthers<sup>18</sup> |
| proROP4 | rabbit | 1:750 |  | Gary Ward<sup>19</sup> |
| ROP2,3,4 | mouse | 1:250 |  | Maryse Lebrun<sup>20</sup> |
| SAG1 (DG52) | mouse | 1:500 |  | Jeroen Saeij<sup>5,21</sup> |
| SAG1 (T41E5) | mouse &<br>rabbit | 1:500 |  | Jean-François Dubremetz<sup>22</sup> |
| SERCA | mouse | 1:5000 |  | David Sibley<sup>23</sup> |
| α-tubulin (12G10) | mouse | 1:2000 | 1:100 | Developmental Studies<br>Hybridoma Bank<sup>24</sup> |
| Tg-β-tubulin | rabbit | 1:500 | | Naomi Morrisette<sup>25</sup> |
| VP1 | rabbit | 1:4000 |  | Silvia Moreno<sup>26</sup> |

